# UC2 – A Versatile and Customizable low-cost 3D-printed Optical Open-Standard for microscopic imaging

**DOI:** 10.1101/2020.03.02.973073

**Authors:** Benedict Diederich, René Lachmann, Swen Carlstedt, Barbora Marsikova, Haoran Wang, Xavier Uwurukundo, Alexander Mosig, Rainer Heintzmann

## Abstract

With UC2 (You-See-Too) we present an inexpensive 3D-printed microscopy toolbox. The system is based on concepts of modular development, rapid-prototyping and all-time accessibility using widely available off-the-shelf optic and electronic components. We aim to democratize microscopy, reduce the reproduction crisis and enhance trust into science by making it available to everyone via an open-access public repository. Due to its versatility the aim is to boost creativity and non-conventional approaches. In this paper, we demonstrate a development cycle from basic blocks to different microscopic techniques. First, we build a bright-field system and stress-test it by observing macrophage cell differentiation, apoptosis and proliferation incubator-enclosed for seven days with automatic focussing to minimize axial drift. We prove versatility by assembling a system using the same components to a fully working fluorescence light-sheet system and acquire a 3D volume of a GFP-expressing living drosophila larvae. Finally, we sketch and demonstrate further possible setups to draw a picture on how the system can be used for reproducible prototyping in scientific research. All design files for replicating the experimental setups are provided via an open-access online repository (https://github.com/bionanoimaging/UC2-GIT) to foster widespread use.

## Introduction

### General

The growing demand in biological research for spatial and temporal resolution, image volume, molecular-tracking and high-throughput coined increasingly complex and expensive light microscope aiming to resolve and track features on molecular level at small time-scales[1, 2]. In order to circumvent the optical resolution limit established by Ernst Abbe [3], these systems need high stability and quality of optical and mechanical components hence resulting in very expensive and complicated setups. Alongside different imaging modalities, long-term observations of living organisms with the least possible bias on their behaviour became an important aspect in microscopy. Keeping the cells in a well-controlled environment and state pose additional constraints like imaging inside an incubator [4, 5] with appropriate microscopes [6, 7, 8] or on-microscope incubator units [9].

Assembling, maintaining and improving such setups as well as analysing and verifying the produced data very often requires a specialist dedicated to the respective instrument, separating microscope engineers from microscope users. Recent approaches like the Flamingo [10, 11] try to enable light-sheet microscopy-as-a-service to everyone thereby resolving an issue of accessibility, but still keeping the gap between developers and users. Despite such rare new approaches, users typically have to stick to already available imaging methods in their lab, therefore preserving traditional ways of data generation and scientific questioning.

The influence of open-source approaches leads to public-accessible workshop- and project documentation. Even though guides on how to build systems like e.g. the lattice light-sheet [12] or openSPIM [13] can be found online, it still needs trained experts for system-assembly. A high price-tag further hinders wide spread and promotes disbelief and a reproducibility crisis in science [14]. Even though optical lab suppliers provide hardware for prototyping, adaption between different distributors and standards (e.g. DIN, ISO, RMS, etc.) often leading to time-consuming handcrafting, an open standard in optics and microscopy is clearly missing.

Analogous to the revolution in biological imaging, the growth of 3D-printing and its price-drop along with inexpensive readily available opto-mechanical consumer components had big impact on existing development paradigms leading to “rapid prototyping” and therewith influencing industrial, academic and home environments [15, 16, 17].

In 2005, the open-source microcontroller Arduino[18] was introduced. Its integrated desktop programming environment and the availability of libraries made microcontroller-development possible for everyone. Together with Raspberry Pi [19] and the ever growing capabilities of smartphones, these technologies push-forward the Internet-of-Things (IoT) applications and change the landscape of education-paradigms e.g. in electronics [20]. Especially smartphones, equipped with mass-produced high-precision optics and tailored image processing algorithms on powerful computational units, can serve as high performance imagers. With over 2.3 billion phones in daily use [21], high quality research or data acquisition can be performed remotely fostering telemedicine applications [22].

### Computational- and 3D-printed Microscopy

Computational microscopy circumvents the possible loss of image quality, caused by the selection of components of lower quality, by combining tailored image processing algorithms with known variations in the physical experiment, like variation of the point-spread functions (PSF)[23] or added hardware like LED-Arrays for quantitative imaging [24]. Thereby, enabling new ways of experimental designs and scientific questioning on the budget.

In terms of hardware design, the monolithic *open-flexure* stage [25] provides a 3D-printed microscope body enabling refocussing of the sample which would be difficult to achieve with former injection-moulding or metalworking technologies. Although being notably stable the system inherently lacked easy adaptability to different imaging configurations. The *100€ lab* [26] relies more on single parts thereby providing easier customizability, but still being quiete specialized. Approaches like the *Foldscope* by [27] and the *cell* STORM [28] show the potential how cellphones can be used for cutting-edge research.

For bio-medically relevant imaging techniques such as fluorescence, a light-weight portable mini-microscope *Miniscope* [29], novel waveguide-based on-chip fluorescent measurement devices [30] and open-source single-molecule localization microscopy (SMLM) system *miCube*. [31] were presented. All systems represent a case-specific prototype, but are hard to adapt to other uses and therefore rather inflexible.

A more generic approach, in the form of an opto-mechanical toolbox, was presented [32] and a functional-unit box-like approach (µCubes) [33]. Although the latter approach already included a functional-unit separation as well as trying to avoid any metallic/pre-manufactured holders, it lacked the freedom of adjustment within the blocks on operation, quick “plug-and-play” change of position of individual parts and crossover compatibility.

With our approach UC2 (pronounce “You, See, Too”) we try to address some of the short-comings and present a modular, easy to build and use optical toolbox equipped with open-source software, design-files, blueprints for a large variety of setups and accessible documentation for the use as innovative tools in labs as well as educational areas. This creates inexpensive microscopic imaging devices for around 100-300 Euro by relying on off-the-shelf components for everyone, anywhere, any-time (Supp. Chapter 3).

### Biological Application

As a first application of our tool, we analyse the *in-vitro* differentiation of monocytes to macrophages. As part of the innate immune system [34], macrophages notably invade their ambient tissue, but also act as scavenging cells and are involved in the clearance of pathogens and dead cells[35]. Blood-born monocytes can be isolated and differentiate within 7 days in the presence of granulocyte macrophage colony stimulating factor (GM-CSF) and macrophage colony stimulating factor (M-CSF) to macrophages. During differentiation process they increase in size [36] and are able to change their morphology depending on their polarization [37, 38, 39]. We demonstrate reproducibility of UC2 experiments by building four autonomous incubator-enclosed bright-field microscopes to observe the differentiation process and possible morphological changes of monocytes *in-vitro*. The experience can then be transferred to any other cell-type for any other imaging based cell assay, e.g. wound healing, cell cytotoxicity, chemoattraction or cell adhesion in microfluidic devices. Finally, we demonstrate versatility and creativity-support of the toolbox by switching to other imaging techniques like selective plane illumination microscopy (SPIM, [13]), quantitative phase imaging (QPI) or structured illumination microscopy (SIM) for example.

## Results

### The UC2 Unit Building-Block

Modern microscopes with infinitely corrected objective lenses often follow the *4f* -configuration, where lenses are aligned such that focal-planes (*f*) of adjacent elements coincide to limit the amount of optical aberrations, to realize telecentricity and to predict the system behaviour using Fourier-optics [40].

To promote the modular property of these systems, we created a generic framework around a simple 3D-printable cube (Fig 1 b). By analysing many available optical components, imaging systems and frameworks we found that a design pitch of *d*_*block*_ = 50 *mm* seems to optimally balance compatibility, handling and flexibility for enabling Fourier optical setups. Separating the cube into a base and a lid simplifies 3D printing using off-the-shelf fused deposition modelling (FDM) printers and allows to insert inner components as plug-ins easily.

**Fig 1.**
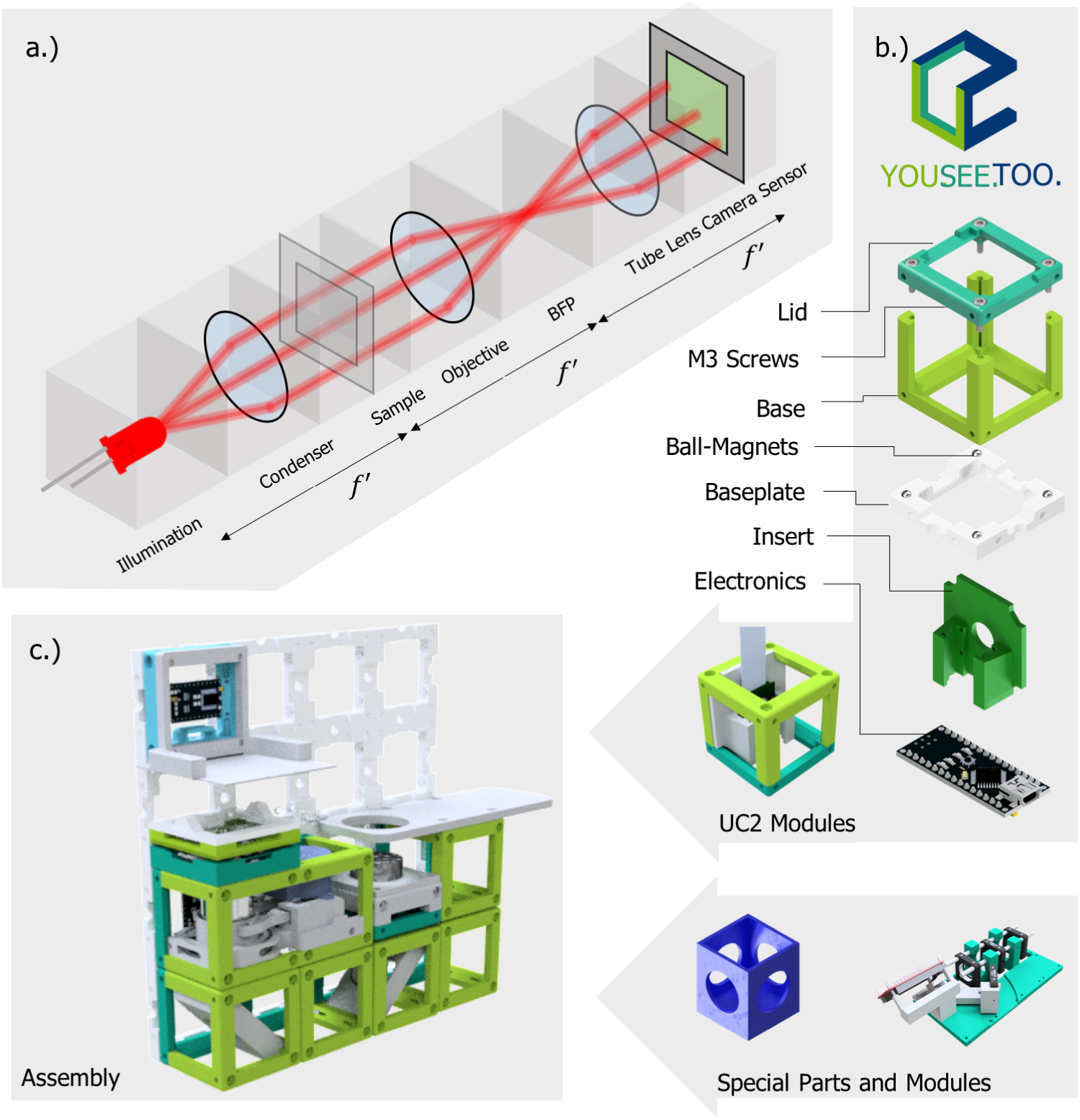
Optical Setup UC2 Blocks. a) The *4f* -system divides Fourier-optical arrangements into functional blocks, where *f*′ corresponds to the focal-lengths. b) The unit element (cube) acts as a base framework for any component which fits inside (lens, camera, z-autofocusing mechanism, etc). b) A magnetic snap-fit mechanism connects the building blocks to a skeleton to realize mechanical stability and rapid-prototyping of a given optical setup. c) An exemplary setup of a microscope for an ordinary smartphone (not shown) and an inexpensive objective as a combination of available modules.

Having neodymium ball magnets (∅_*magnet*_ = 5 *mm*) rectangularly oriented on an extendable base-plate and ferro-magnetic cylindrical bolt screws (DIN 912) sitting in the cube’s edges allows a stable and precise magnetic mount in three dimensions. We found a four-point fixation as a good compromise between the common rectangular arrangement of optical setups and mechanical stability.

External electro and optical components (Raspberry Pi-camera, Mirrors, LEDs; see Fig 1 b) and already existing equipment (e.g. rail-systems from Thorlabs, Quioptics, Edmund Optics, etc.) can be easily adapted by plug- and modifiable inserts. A module developer kit (MDK, Supp. Chapter 3) with a generic reference design for customized inserts provides a simple interface for even non-technical users to work or add designs to the toolbox.

### From Modules to Assemblies

Scaling complexity of optical systems starting from a simple magnifying glass up to a fully-working light-sheet setup (Fig 2) is ensured by relying on the previously introduced library of modules which are combined and put in the appropriate order (Supp. Chapter 3). Adding more advanced consumer electronics (cameras, motors, video-projects, etc.) allows the use as smart-microscopes and enable remote control. Micro-controllers ensure wired (i.e. *I*^2^*C* [41]) or wireless (i.e. WiFi, IoT-based protocol using Message Queuing Telemetry Transport (MQTT) [42]) communication interface to trigger light-settings or focussing mechanisms Supp. Chapter 3). Power is supplied through the conducting magnets.

**Fig 2.**
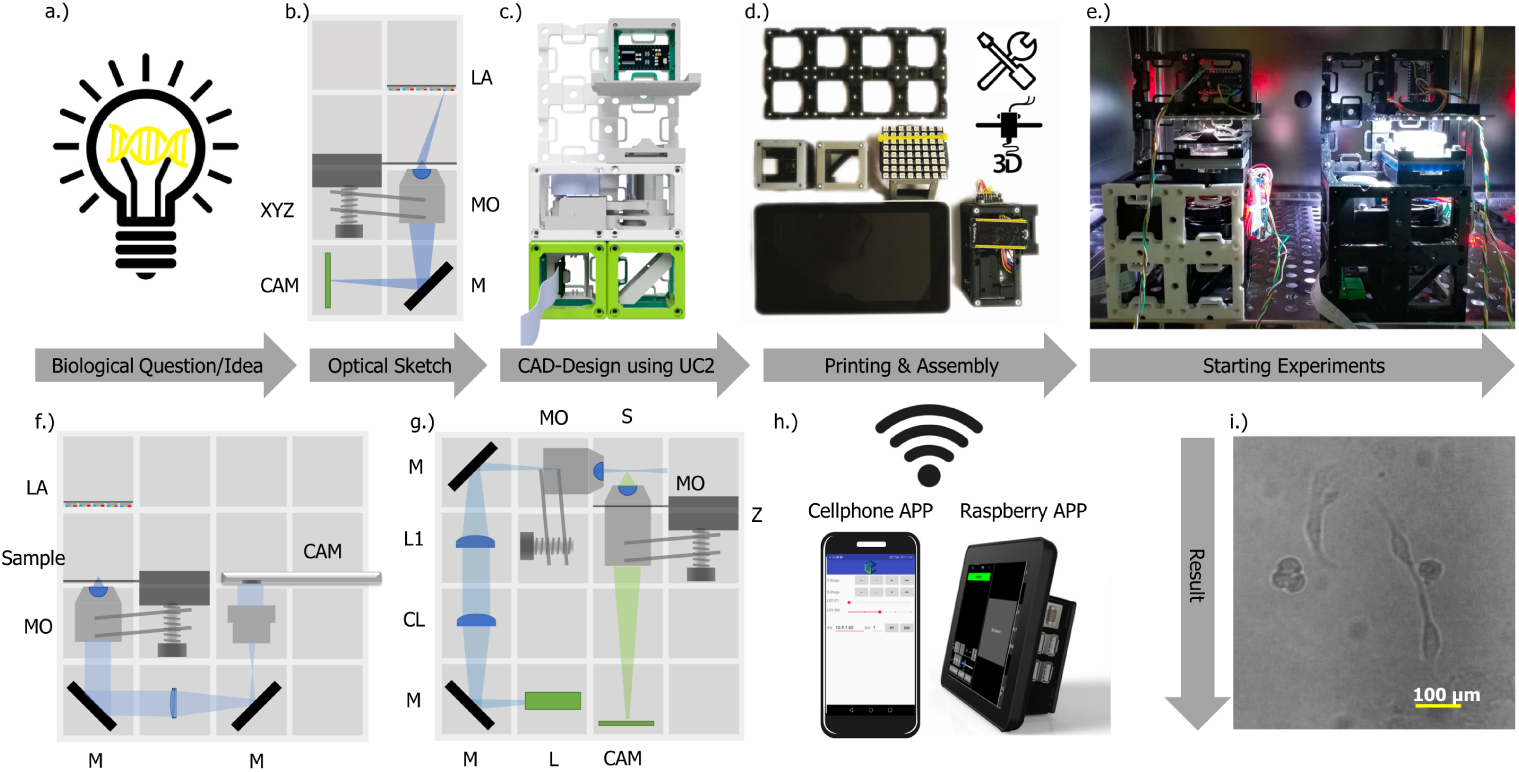
Rapid Prototyping using UC2. - Common workflow to build a UC2 application: a) Starting with a biological question/idea in need of an imaging device drafted in b) (inverted incubator microscope) and transferred using UC2 components from the CAD library in c). After printing and assembling it d) the device will be placed in its working environment (e.g. incubator) e) ready to acquire long-term image series visualized in i). Remote-controlled is granted using MQTT-enabled devices (cellphone, Raspberry Pi) in h). Reusing components allows the conversion into a cellphone-microscope f) or light-sheet microscope g) within minutes (see Supp. Video 2). *CL*: cylindrical lens, *L*: Laser, *LA*: LED-array, *L1* : Lens 1, *M* : Mirror, *MO* : Microscope Objective, *P* -CAM: Detector (Smartphone or Raspberry Pi), *S* : sample positioning stage, *XYZ* : axial- and lateral translation stage, *Z* : axial translation stage

### Configuration 1: Compact Device for Long-term *in-vitro* Imaging

To minimize environmental effects such as infection of the cells and to democratize access to *in-vitro* imaging tools, we built a small inverted microscope (Fig. 2) in bright-field-mode (BF) with an optical resolution on cellular-level (i.e. < 2.2 *µm*) for ≈ 300 Euro (Fig. 2 b)-e)). For cross-verification, stability measurements and display of parallelization we placed four BF-setups (2× *I*^2^*C*-, 2× MQTT-interface) into a single incubator. We specifically designed a graphical user-interface (GUI) on the Raspberry Pi, to preview the region-of-interest, set the imaging parameters (focus, illumination) and ensure autonomous image acquisition (Supp. Chapter 3). We performed multiple long-term measurements under conditions of high humidity (≈ 100%) and at temperatures around 37°*C, CO*_2_ = 5% over 7 days taking images with 1 frame per minute repetition rate thereby continuously monitoring the morphological changes and plasticity related monocyte-differentiation. We placed isolated monocytes in 35 *mm* dishes rinsed with 3 *ml* X-Vivo (Lonza, GA, USA). The shape of the differentiating monocytes is round and elongates while moving (not further quantified). The increase in area is also shown in Fig. 3 d) where the mean-area of individual cells over a subset of time-points is plotted. We observed a significant increase of size by 4 times within 100 *min* observation as verified by average-control using GraphPad Prism (ANOVA with post-hoc Turkey’s; CA, USA). Macrophages are used to “discover” their surrounding by extracting their plasma membrane in pseudopodia in order to detect pathogens or cell debris. We even detected an apoptotic macrophage being phagocytosed by the surrounding macrophages (Supp. Fig. 1) and rarely seen division of a macrophage (Supp. Fig. 2).

**Fig 3.**
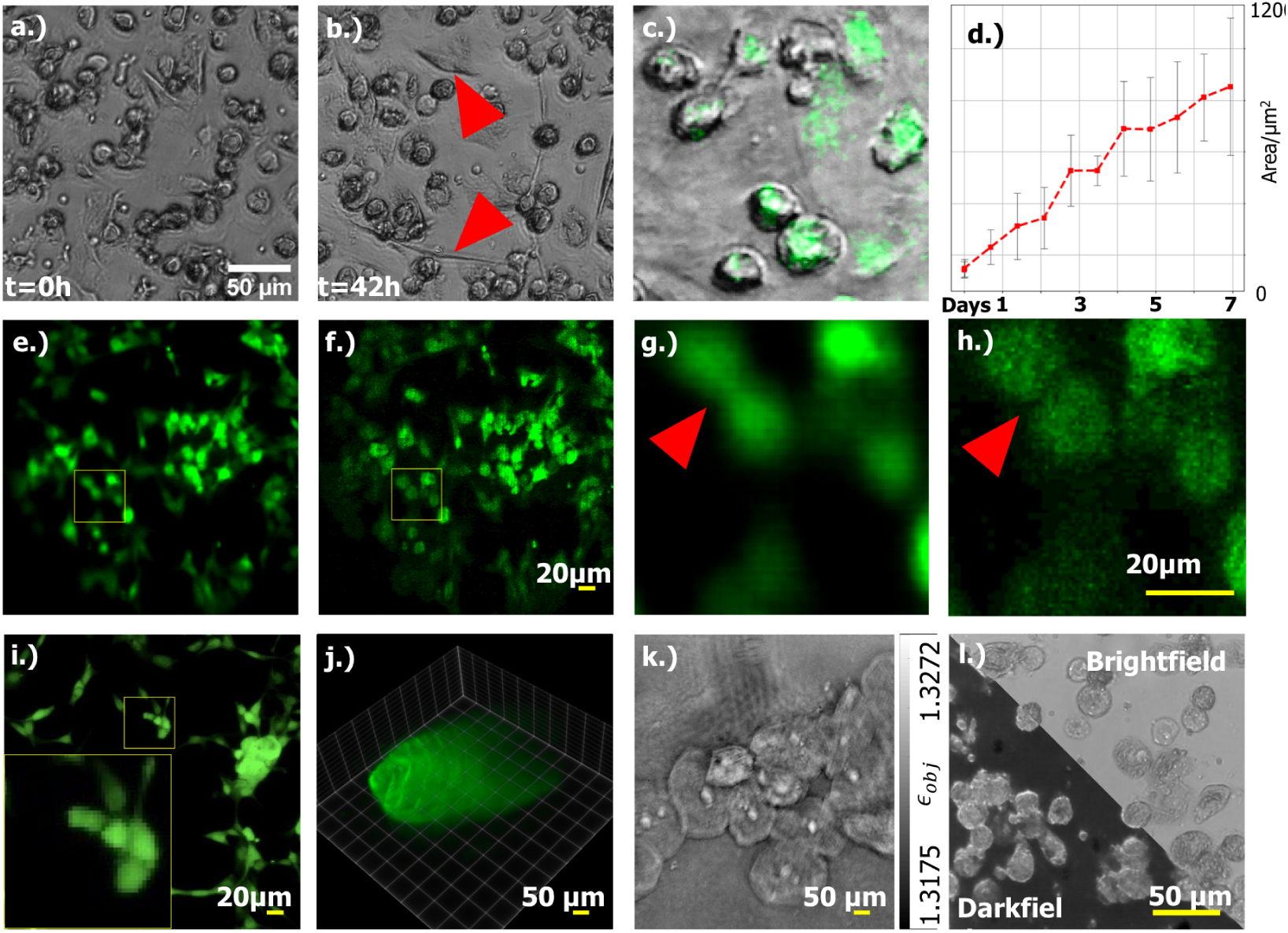
Visualizing the different imaging modalities from a UC2 setup. Variation in macrophage’s morphology In a)-b), where elongated cells are clearly visible after 42 *h* (red arrow). The growth of a differentiating cell is plotted as the average area of cells across multiple time-steps and different experiments in d). c) The bright-field channel superposed with a fluorescent signal of fixed macrophages labelled with CellTracker green captured with the incubator-enclosed microscope. e) Wide-field fluorescence and f.) the computed “superconfocal” result of GFP-labelled HPMECs, illuminated with a laser-scanning projector, recorded with a cellphone camera. The zoomed-in images show the improvement of the optical sectioning in case of structured illumination in h.) compared to widefield in g.), where smaller cell-structures are lost. i) A comparison of the same sample acquired with a commercial laser-scanning confocal microscope. j) An acquired z-stack of a GFP-expressing drosophila larva. k.) Using an LED-ring as the illumination enables quantitative phase imaging of cheek cells using *A-IDT*. l) LED matrices can rapidly switch between bright- and darkfield imaging as shown in l)

During imaging, the magnetically-mechanically fixed *in-vitro* sample (e.g. ∅ = 35 *mm* petri-dish, organ-on-a-chip or standard microfluidic chips e.g. Ibidi µ-chip) experiences a focus drift due to temperature-dependent bending (See Supp. Video 2) which was compensated by software autofocus. A later introduced spiral flexure-bearing-based stage printed using Acrylnitril Butadien Styrol (ABS) improved the stability resulting in negligible focus-drift after the pre-warming phase.

Fig. 3 c) shows an exemplary overlay of fluorescent- and bright-field signal using the back-illuminated colour CMOS sensor of a Raspberry Pi camera of fluorescently labelled (CellTracker Green, Thermo Fisher) monocytes following in a rather low signal-to-noise ratio (SNR). An improvement is obtained using monochromatic back-lit CMOS sensors from a cellphone camera (e.g. Huawei P20, China) which can be used by adding an eyepiece to get a correct imaging condition (Fig. 2 g.). Additionally, to realize image scanning microscopy (ISM), [43], Supp. Chapter 3), we added a customized module hosting a laser-scanning video-projector (Sony MP.CL1A, Japan) with peak wavelength of *λ*_*blue*_ = 450 *nm* to excite GFP-labelled HPMEC cells. Combined with an appropriate filter cube we measured fluorescent images in an infinity corrected (Optika, N-Plan, 20×, *NA* = 0.4, 80 Euro, Italy) setup (Supp. Chapter 3) comparable to a state-of-the art laser-scanning confocal microscope (Leica TCS SP5, Fluotar 20×, *NA* = 0.5, Germany) visualized in Fig. 3 e-i). The computationally reconstructed “superconfocal” image [44] Fig. 2 h.) shows better optical sectioning compared to the wide-field equivalent g.).

Using the same setup, further imaging modalities can be easily applied and tested. Due to using an LED matrix (Adafruit #1487, NY, USA) as light-source (in transmission mode) the selection of the illumination wavelength, particular pattern for contrast-maximization [45], dark-field illumination or quantitative phase-methods like “(quantitative) differential phase contrast” (*q* DPC, [24], see Supp. Chapter 3) and “Fourier Ptychography Microscopy” (FPM, [46]) are easily possible. We changed the matrix with a LED ring (Adafruit#1463) to demonstrate computational refocussing of a recovered phase map of cheek cells (Fig. 3 k)) by applying the “Annular Intensity Diffraction Tomography” (aIDT, [47], see also Supp. Chapter 3) method. We implement fluorescent imaging via high-power LEDs of different wavelength in a dark-field configuration from below the sample to reduce cross talk caused by spurious non-fluorescent stray light [48].

### Configuration 2: Light-sheet Microscope for Educational Areas

By adding a small number of components it is possible to reconfigure the previous incubator-enclosed system into a light-sheet microscope inspired by *open*SPIM [13] within minutes (Supp. Chapter 3, Supp. Video 2). We acquired a 3D data-stack of zebra-fish embryo and drosophila larva expressing GFP (Fig. 3j, Supp. Fig. 3). The acquired data was drift-corrected and deconvolved to achieve a reconstructed resolution.

Finally, we analysed the minimum necessary amount of printed and off-the-shelf components to build the formerly mentioned setups as well as telescopes, projectors, Abbe-diffraction experiments or holographic (e.g. lens-less) imaging devices and compiled a ready-to-print collection of open-sourced parts and documentation - named *TheBOX* (see Supp. Chapter 3) and a version optimized for microscope courses *CoreBOX*. It is supported by a continuously improving documentation with step-by-step guides and tutorials. We tested the system at various conferences, workshops and educational environments, and obtained feedback loop to further improve the system.

## Discussion

Born from the idea to stop reinventing the wheel when creating a new microscopy method we introduced a new modular toolbox that we belief has the potential to serve as the new truly open-standard, not only in microscopy. Its inherent availability and ease-of-use already enabled many people - from school pupils over private enthusiast to researchers - to build and work with their own systems. With the application of macrophages long-term imaging presented here, we addressed the simplification and barrier reduction into optical research thereby inviting curious minds from different backgrounds to interact with, find novel methods of data-acquisition or processing or to verify and test new microscopic methods. The system cultivates the inherent “Spieltrieb” (play instinct) of humans by reducing fear of potentially destroying expensive components or long setup construction.

We proved reproducibility not only externally, but also by building four incubator-enclosed systems and long-term testing them in one incubator with *in-vitro* macrophage differentiation, further proving the benefits of its inherent small footprint. The macrophages in our setup showed the expected increase of size [36]. We were able to distinguish the macrophage body from the pseudopodia [49] and to follow their movements and track their shape. The untreated macrophages show in the majority a roundish shape that gives hint on their not activated phenotype. Nevertheless inactivated macrophages are assumed to be fusiformed [39]. Having access to long-term acquisition enables to relate the elongated form of the macrophage to its movement which is increased in an activated macrophage population [50].

With this established and reliable microscope for a very low budged, high trough put and customized applications like microfluidic chips can be realized.

Further, we showed versatility and flexibility by demonstrating different image-modalities with a) the same incubator-enclosed configuration and b) a SPIM-like system. Even though the system is limited in robustness and stability, we were able to image a living drosophila larva. Taking the low price-tag into account, being able to dispose of contaminated microscopic-systems in high-safety biological environments is now possible. It also serves as an affordable tool for Ebola outbreaks.

The system is highly modular on the hardware as well as on the software-side allowing for easy integration into existing systems, like the Openflexure stage [25], µCube [33], Micro-manager [51] and ImJoy[52]. Furthermore, the existing pool of ready-to-use modules enables rapid prototyping not only in biology, but also in algorithm development or educational environments. With this, the UC2 system strives to fill the gap of what the Arduino is for electronics and Fiji for (microscopy) image processing. Although, the UC2 system is not (yet) intended to produce highest quality images, it nevertheless is a toolbox to create comprehensible experiments. Even though we clearly defined spatial dimensions and used materials, the provided toolbox can easily be rescaled and produced in different materials. As an example, by shrinking and closing the cubes to optimally work with available ∅_*lens*_ = 2 *mm* lenses, even in-vivo medical systems or small-scale microfluidic systems might be worth thinking about.

We introduced a sophisticated tool set for educational purposes with *TheBox*. The compilation costs around 600 Euro including a monitor-equipped compter and is of similar quality as commercial instruments with one to two orders of magnitude higher price tags. Together with a series of ready-to-use documentations optical concepts (e.g interference, image formation, etc.) and basic to complex microscopy methods can be visually demonstrated. This gives students and users (e.g. at Universities, High-schools, extracurricular educational centres, imaging facilities, etc.) the possibility to experience how optics works by trying it themselves and directly addresses the crisis found in STEM (Science, Technology, Engineering and Mathematics) education by promoting interdisciplinary approaches where several educational topics are treated at once. Exemplary teaching material is given in Supp. Figure 3.

With the UC2 system we presented an eager approach to democratize research and make state-of-the-art techniques available to everyone. We try to counter the reproduction-crisis and restore belief in natural sciences by opening new ways of peer-reviewed research, where protocols now also apply to hardware and experiments can be retraced directly. This way more advanced imaging schemes (e.g. ISM in Supp. Chapter 3, SIM in Supp. Chapter 3) or prospective methods can be created by incorporating and sharing the expertise of individual research groups. We hope that this system in combination with an intuitive assembly and usage-description gets adopted by the research community to react on individual and global needs in microscopy. The development process can be split to a larger number of people, where participation and new influences can improve the result dramatically. Thereby the system will benefit from a wide variety of contributors. The function as an imaging device thus no longer results at the moment of the production but rather from the sum of the individual parts in combination with the creativity of the user and allows new ways of data creation e.g. for machine or deep learning to restore reproducibility on a global scale.

## Methods

### Fabrication of the components and selection of additional parts for the Incubator-Microscope

A detailed description of each individual part as well as the bill of material (BOM) can be found in our Supp. Material available in the GitHub-repository at http://github.com/bionanoimaging/UC2-GIT (Supp. Chapter 3). In general, all components of the UC2 toolbox are designed using common CAD software (Autodesk Inventor 2019, MA, USA) and were printed using off-the-shelf FDM-based 3D printers (Prusa i3, mkII, Czech Republic; Ultimaker 2+/3, Netherlands), where in most cases poly-lactic acid (PLA, *t*_*print*_ = 215°*C*) was used as the material. The infill was chosen between 20%-40% together with a layer-height of 0.15 *mm* which provides high enough precision and stability for all optical setups. The monolithically printed z-stage cube (Supp. Chapter 3) based on a linear or spiral flexure-bearing and certain base-plates for the use in the incubator were printed using ABS which provided better long-term stability at *t*_*incubator*_ = 37°. It adapts to common objective lenses (e.g. RMS-thread) which gets linearly translated using a worm-drive realized with a M3 screw and nut driven by an inexpensive stepper motor (28BYJ-48, China).

Black material was used in most cases to reduce stray light or unwanted reflection and scattering. To decontaminate the printed parts, the assembled cubes were sprayed with 70% ethanol before entering the live-cell imaging lab facility.

For the magnetic snap-fit mechanism 5 *mm* neodymium ball-magnets were press-fit into the printed base-plate which adapted to M3×18mm galvanized cylindrical screws (Würth M3×18, ISO 4762 / DIN 912) sitting in each face of the cube to assure a stable connection. Additional wires added to the magnets and screws respectively can support electro-optical modules (e.g. LED array) with electrical power (i.e. 5V, GND), where rectifier can prevent wrong polarity. To keep the optical design simple and compact we relied on a low-cost (15 Euro) finite corrected objective lens (10×, NA=0.3, China), where the beam was folded using a cosmetic mirror (20 cents). The image formed at a reduced tube-length *d*_*tube*_ = 100 *mm*) was captured using a back-illuminated CMOS sensor (Raspberry Pi Camera, v2.1, UK), connected to a Raspberry Pi v3. An additional module which incorporates a pair of motor-driven, low-cost XY micro-stages (3 Euro, *d*_*x*_ = *d*_*y*_ = ±1.2 *mm*, Aliexpress, China) to place the sample precisely in XY (Supp. Chapter 3). For bright-field and quantitative imaging we used a 8 × 8 LED-array (Adafruit #1487, NY, USA) where a GUI was run on a 7-inch touchscreen (Raspberry Pi, UK) provides selection of individual LEDs to maximize the contrast according to Siedentopf’s principle [45]. For fluorescent imaging of GFP labelled HPMEC cells, we equipped the Fluorescent-module (Supp. Chapter 3) with high-power star-LEDs (Cree, 450nm/405nm+/-20nm) and added a gel colour filter in front of the CMOS sensor (ROSCO #11).

### Hardware Synchronisation and Image Acquisition

All sources together with a full documentation of the software briefly described below together with an in-detail set of instructions can be found in our GitHub repository and Supp. Chapter 3. A reduction of wires for “active” modules (e.g equipped with motors, LED’s, etc.) was achieved by a microcontroller connecting to a wired *I*^2^*C*-BUS (Arduino Nano, Italy) or a wireless MQTT protocol based (ESP32 WROOM, China) network. As a master device for the 4-wired *I*^2^*C* connection we choose the Raspberry Pi v3B, using *I*^2^*C*. The ESP32 can be controlled with any MQTT-device, e.g. Raspberry Pi, Smartphone or other ESP32/Arduino microcontrollers in the same network making it also possible to control the device remotely (e.g. from the office).

A user-friendly Python-based [53] GUI running on a 7-inch touchscreen gives access to functions like scheduling experiments, setting up imaging modalities (e.g. illumination pattern) and hardware-/frame synchronization for several applications (e.g. incubator-enclosed microscope). Frames from the camera module (Raspberry Pi, v2.1) are stored as compressed JPEG images to save memory. In cases where cellphones (e.g. Huawei P9/P20, China) as imaging devices were used, the open-source camera APP FreeDCam ([54]) was used to have full control over imaging parameters (i.e. ISO, exposure time, etc.) and access to RAW images. USB-batteries (e.g. power banks) allow autonomous operation in rural areas over several days.

We tested live-drift-correction to account for vibration or expanding of the material by a software-based autofocus (i.e. axial defocus), where we used a direct spatial filter (i.e. Tennengrad) [55] as an image sharpness-metric. This autofocus was not used for final measurements due to the inherent stability of the new z-stage.

### Image Analysis and Image Processing

A customized Python [53] script handles long-term measurements (e.g one frame-per-minute over 1 week) by binning the RAW-data and creating a preview video. Then manually a frame of reference, where the lateral sample-drift seems to have settled, and regions-of-interest (ROI) for fix image features - here: dirt on sensor - were defined. On a second iteration, image statistics like min,max,mean or image-sharpness and shift, using a cross-correlation estimation, - for the whole image and the ROIs - with respect to the reference frame are calculated. Due to the large temperature coefficient (1.01 × 10^−4^*K*^−1^) the ABS tends to deform especially dominant during the one-hour heat-up phase in the incubator. Finally, dark and corrupted frames are excluded using the statistical measures, shifts are applied and a stack-mean is calculated for the green channel. Finally, by only the green channel is further processed. Flat-fielding and dirt-correction is achieved by division through the stack mean. Finally, division by per-frame mean and normalization to the same stack mean is done to account for unequal illumination and sensor-errors (e.g. dirt, scratches).

For task-specific image processing on the cellphone directly such as the processing of the ISM measurements or frame segmentation, we used the cloud-based image processing framework ImJoy [52] for available on our GitHub repository (Supp. Chapter 3).

For the quantitative phase measurements based on the aIDT, we used the publicly available Matlab (The MathWorks, MA, USA) code from Li et al. [47] with small modifications according to the optical system using the cellphone microscope (see Supp. Chapter 3).

Possible fluctuation of Z-stacks acquired with the light-sheet microscope were registered using a cross-correlation based routine before a blind-deconvolution based on the publicly available *DeconvToolbox* by Heintzman et al. removed out-of-focus blur. Fiji [56] was used for measuring the cell size (i.e. macrophages), the diameter was determined manually across 10 time-frames over the whole 1-week measurement. In each frame, an individual cell was selected manually before the round-factor was computed using a customized macro.

### Sample Preparation

Periferal blood mononuclear cells (PBMCs) were isolated from healthy volunteer adult donors by Ficoll density centrifugation. The study and experimental protocols used therein were approved by the ethics committee of the University Hospital Jena (assigned study number 2018-1052-BO). In detail, the blood was mixed with isobuffer (PBS without Ca/Mg (Gibco), 2 mM EDTA (Sigma-Aldrich, USA), 0.1% BSA (Sigma Aldrich, Germany)) and placed on top of Biocoll (Biochrom, Merck, Germany) without mixing in a 50 *ml* tube. Biocoll and Blood were centrifuged with 800 x g for 20 min with out break. The ring of PBMCs was transferred in a new 50 *ml* tube and washed twice with isobuffer. PBMCs were seeded at a density of 1 *x* 10^6^*cells/cm*^2^ in X-VIVO 15 medium (Lonza, Cologne, Germany) supplemented with 10% (v/v) autologous human serum, 10 *ng/ml* granulocyte macrophage colony stimulating factor (GM-CSF) and 10 *ng/ml* macrophage colony stimulating factor (M-CSF) (PeproTech, Hamburg, Germany) and Pen/Strep (Sigma Aldrich). After 1 h PBMCs were washed twice with RPMI and remaining monocytes were then rinsed with X-Vivo with supplements. 16 h after isolation monocytes were washed with prewarmed PBS (*w/o Ca/mg*) and incubated 7 min with prewarmed with 4 mg/ml lidocain and 1 mM EDTA. Detached monocytes were places in a 15 *ml* tube and centrifuged 7 min by 350*xg*. Sediment monocytes were counted and 1.5*x*10^5^ were seeded in a 35 *mm* dish and rinsed with 3 *ml* X-Vivo 15 with supplements. After 24 h were the cells washed once with X-Vivo 15 and the monocytes were rinsed with 3 ml fresh X-Vivo 15 with supplements and placed in the microscope.

## Authors Contribution

**Conceptualization:** B.D., R.L., S.C.

**Data curation:** B.D., R.L., S.C., B.M., H.W.

**Formal analysis:** B.D., R.L., S.C.

**Funding acquisition:** B.D., R.L., A.M.

**Investigation:** B.D., R.L., S.C.

**Methodology:** B.D., R.L., S.C.

**Project administration:** B.D., R.L.

**Resources:** B.D., R.L., S.C., A.M.

**Software:** B.D., R.L., X.U.

**Supervision:** B.D., R.L., R.H.

**Validation:** B.D., R.L., S.C., B.M., R.H.

**Visualization:** B.D., R.L.

**Writing – original draft:** B.D., R.L., S.C. with input from all authors.

**Writing – review & editing:** B.D., R.L., B.M., R.H.

## Acknowledgments

This study was supported by the Center for Sepsis Control and Care (Federal Ministry of Education and Research (BMBF), Germany, FKZ 01EO1502) and the Leibniz ScienceCampus InfectoOptics Jena, which is financed by the funding line Strategic Networking of the Leibniz Association. Additionally, this work was financially supported by the Deutsche Forschungsge-meinschaft through the Cluster of Excellence “Balance of the Microverse” under Germany’s Excellence Strategy – EXC 2051 – Project-ID 690 390713860 and by the European Commission through Marie Sklodowska-Curie Actions (MSCA) Innovative Training Network EUROoC (grant no. 812954) to A.S.M. The authors want to thank the Lichtwerkstatt Jena – Open Photonics Makerspace located at the Friedrich Schiller University Jena for sharing resources and facilities for multiple workshops. We thank The Leibniz IPHT Jena e.V. for funding the project with the Innovation-fund. Human pulmonary microvascular endothelial cells transfected with eGFP (HPMEC-eGFP) were kindly provided by Dr. Lothar Koch and Andrea Deiwick of the Institute of Quantum Optics, Leibniz University Hannover. We thank Nora Mosig, Melanie Ulrich and Tobias Vogt for their excellent technical assistance. We thank Kaspar Podgorski for hosting and the HHMI Janelia for funding the UC2 workshop at HHMI Janelia Research Farms. We also thank Xian Hi (Edna), Kay Schink and Oddmund Bakke for organising, funding, hosting and preparing drosophila and MDCK samples for the workshop at Oslo University. We thank Philipp Kahn for creating the UC2 project webpage and WiTeLo Jena e.V. for hosting several UC2 workshops. We thank Ronny Förster, Nico Schramma and Kyriacos Leptos for fruitful discussions. Further thanks go to Muriel Starke and Claudia Lachmann for constant support and helpful advises. This work was supported by the GSSP from DAAD.

