## Supplementary Notes 1 for "UC2 – A Versatile and Customizable low-cost 3D-printed Optical Open-Standard for microscopic imaging"

#### UC2 – General Purpose, Low-Cost and Open-Source 3D-Printed Optical Toolbox

#### 1 Supplementary Figure

##### Supplementary Figure 1.

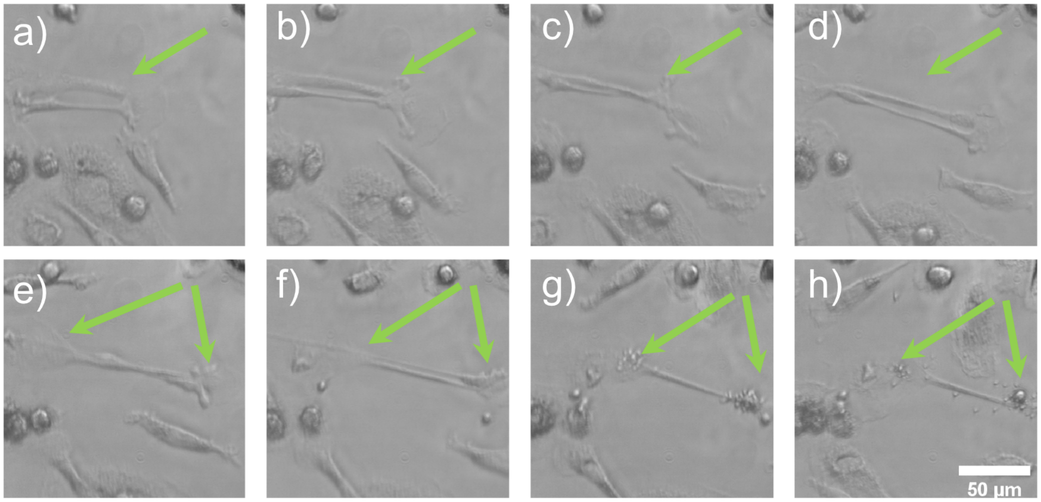

**Fig 1. Cell-Apoptosis of macrophages** From image series with the incubator microscope ($10\times$, 0.32 NA objective) at a frame-rate of 1 frame/minute. a)-d)One observed cell first became elongated and e)-f) started blabbing, a clear sign of apoptosis. g)-h) The fragmentation of the cell to apoptotic bodies is clearly visible. The cell fragments are then cleared via efferocytosis by other macrophages (see Supp. Video 2 from 3:10, left). The displayed images are temporally spaced by 50min each.

**Supplementary Figure 2.**

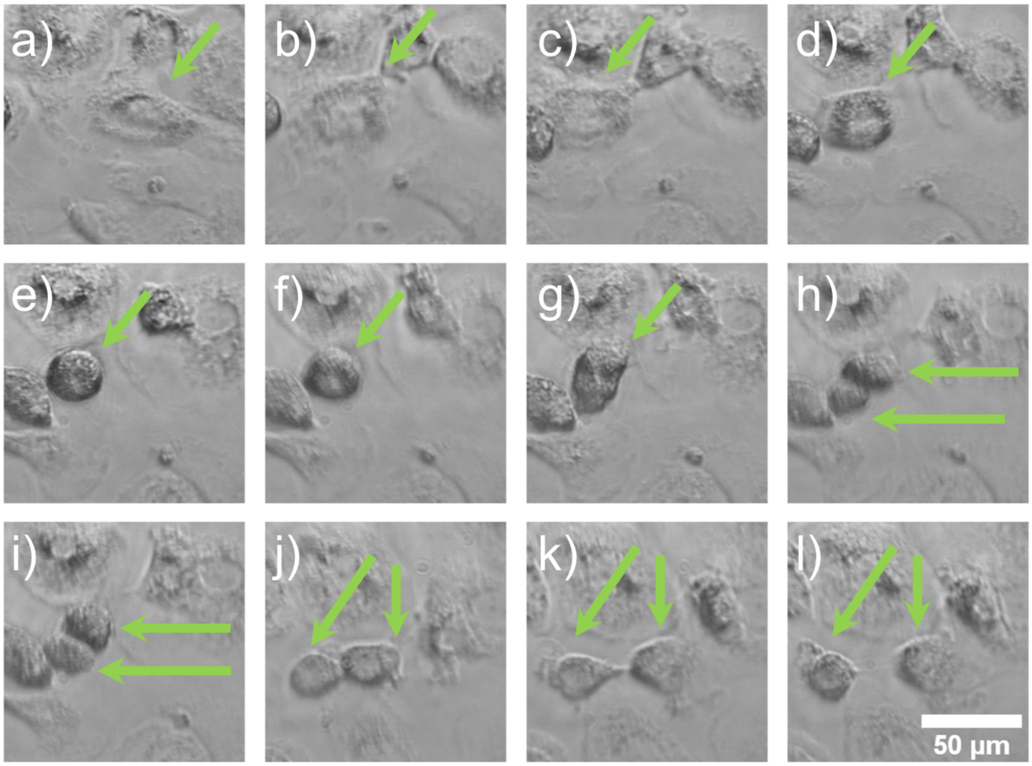

**Fig 2. Cell Division of Macrophages** - Long-term ($48\,h$) image series with the incubator microscope ($10\times$, 0.32 NA objective) at a frame-rate of 1 frame/minute. A very rare cell division of a macrophage was observed. a)-d) Out of a movement the cell stopped, e)-f) constricted and g)-l) divided into two cells. Measurement time-points in additive minutes (min) from starting point a) = 5days 18hours 33min are: 75min, 115min, 129min, 172min, 177min, 182min, 187min, 192min, 748min and 760min. (see Supp. Video 2 from 7:04 - 7:08, center right)

#### Supplementary Figure 3.

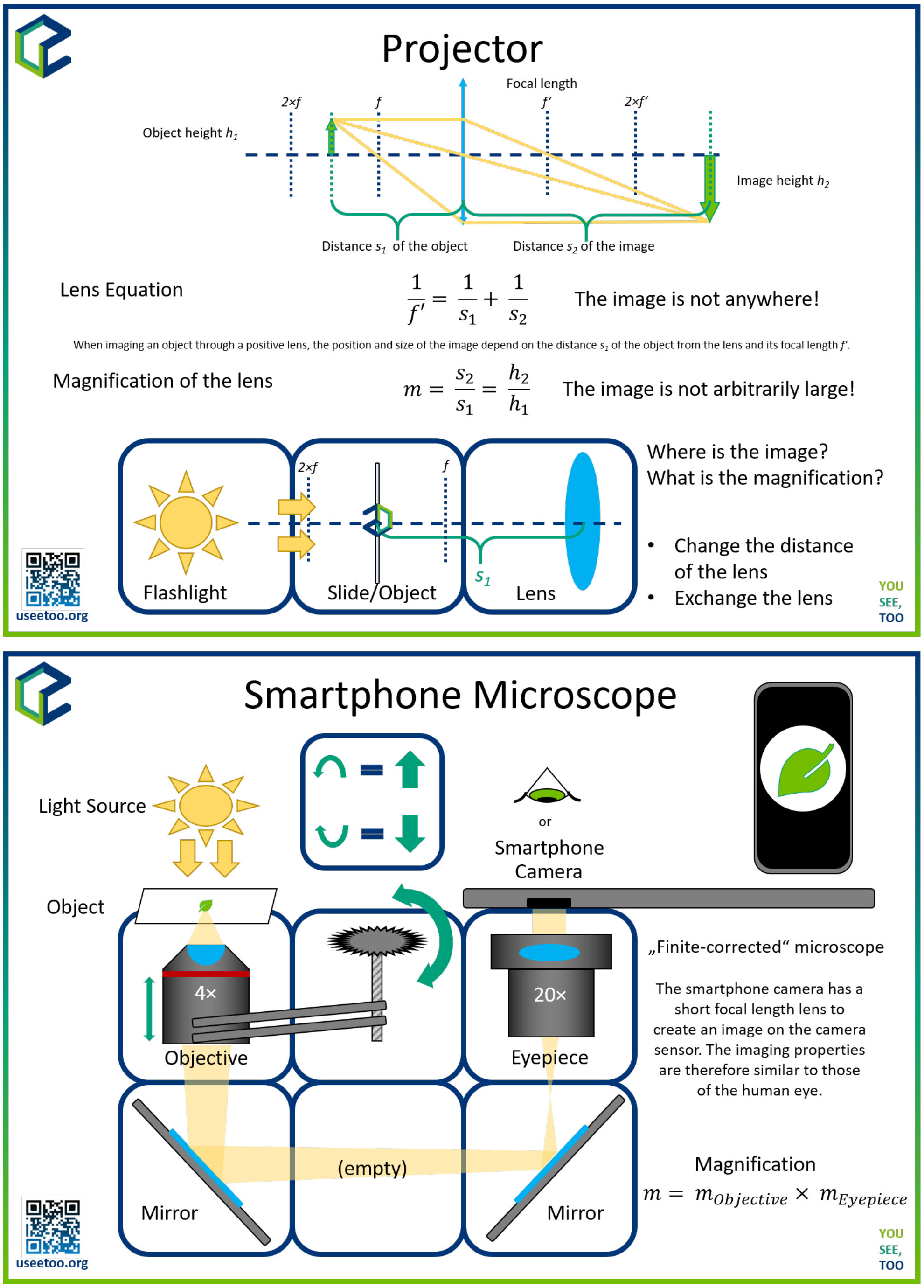

**Fig 3. Educational chart** Exemplary slides to show the basic properties of a simple projector or a smartphone-based microscope for the use in schools. The optical layout acts as a printed template for the different cubes. Students can conveniently place the cubes in these place-holders to create the microscope and observe an image using their eyes or cellphones and discuss the results.

#### 2 Supplementary Video

**Supplementary Video 1.** Long-term ($48\,h$) image series with the incubator microscope ($10\,\times$, $0.32\,\mathrm{NA}$ objective) at a frame-rate of 1 frame/minute. In-Incubator measurement of isolated human blood monocytes. The aim was to document differentiation of Monocytes to Macrophages and analyse their movement pattern without stimulation.

**Supplementary Video 2.** The video shows long-term imaging of the differentiation of blood-born monocytes to macrophages. Within this time span the monocytes increase size and are "looking" around. Obvious are the filopodia round the cells. Moving macrophages become fusiform, elongate and follow their protrusions with the cell body.

**Supplementary Video 3.** Reconstruction of the complex refractive index of unlabelled cheek cells using the annular intensity diffraction tomography algorithm (aIDT). A number of LEDs on a LED-ring placed close to the rim of the back-focal plane of the microscope objective illuminate the sample sequentially, where the inverse filtering process can reconstruct a 3D stack of the permitivity distribution. The acquisition was performed using a cellphone camera (Huawei P20, Pro, China) further described in Chapter 3.

**Supplementary Video 4.** Through-focus series of a Drosophila larva. Due to the large depth of field of the $4\times$, NA=0.17 objective, a data stack can be acquired by moving the light sheet through a fixed sample i.e. by moving the kinematic mirror perpendicular to the detection path. The GFP-expressing drosophila larva was focussed by the detection path and the illumination plane was then moved through it by changing the tilt of the kinematic mirror. Although the whole three-dimensional sample is in focus, only the illuminated parts are imaged onto the camera. The video was acquired with a cellphone camera (Huawei P20, Pro, China).

Alternatively, one can move the whole sample through the fixed light sheet aligned to the focus-plane for the detecting objective lens ($10\times$, NA=0.32) using the sample-stage equipped with a flexure bearing. This is shown in the video of the GFP-expressing zebrafish embryo. The video was acquired with a Raspberry Pi camera.

**Supplementary Video 5.** The conversion from a simple bright field into a light sheet microscope can be accomplished within less than five minutes using TheBOX. The modules can easily be reused for different imaging modalities.The components are pre-aligned and remain their position when packed again, useful for transporting the whole system.

**Supplementary Video 6.** Long-term measurements of MDCK-cells at room-temperature over-night ($8\,h$) in an $35\,mm$ petri-dish at a UC2 workshop given in Oslo, where participants were able to bring their samples.

### 3 Supplementary Notes

#### Module Developer Kit

One aspect which is missing in many open-source and open-science projects is the ability to interact with the project in order to introduce own modifications to individual needs. During our study we found that one major requirement in order to give users easy access to the resources and to make it attractive to start developing on an open project - like the proposed UC2 system - is an easy to understand documentation. It should provide an intuitive way into the project to reduce the inhibition threshold to start working on it.
Inspired by the recently discontinued modular cellphone project *ARA* by Google Inc. [6] we created a comprehensive document called the Module Developer Kit (MDK, GitHub repository) which describes the good practice of cube-design and customized inserts. This includes the CAD-files for common CAD software like Autodesk Inventor 2019 (Autodesk ®Inventor LT™) OpenSCAD (`www.openscad.org`) as well as schematics to port the design to other software tools. It emphasizes the idea of having the UC2-system as a supporting base-structure or skeleton to become a common standard for a large variety of different components of different manufactures. By having a zoo of modules developed by an active community which are useful for many people guarantees a long lifetime of the project.
All files can be found in our hard- and software repository [2, 8].

At first we want to introduce the naming-convention of the UC2 system to give a better understanding of the module hierarchy. These terms are defined in the table (3) below and illustrated in Fig. 4. Based on these modules and inserts, a complex optical system can be created.

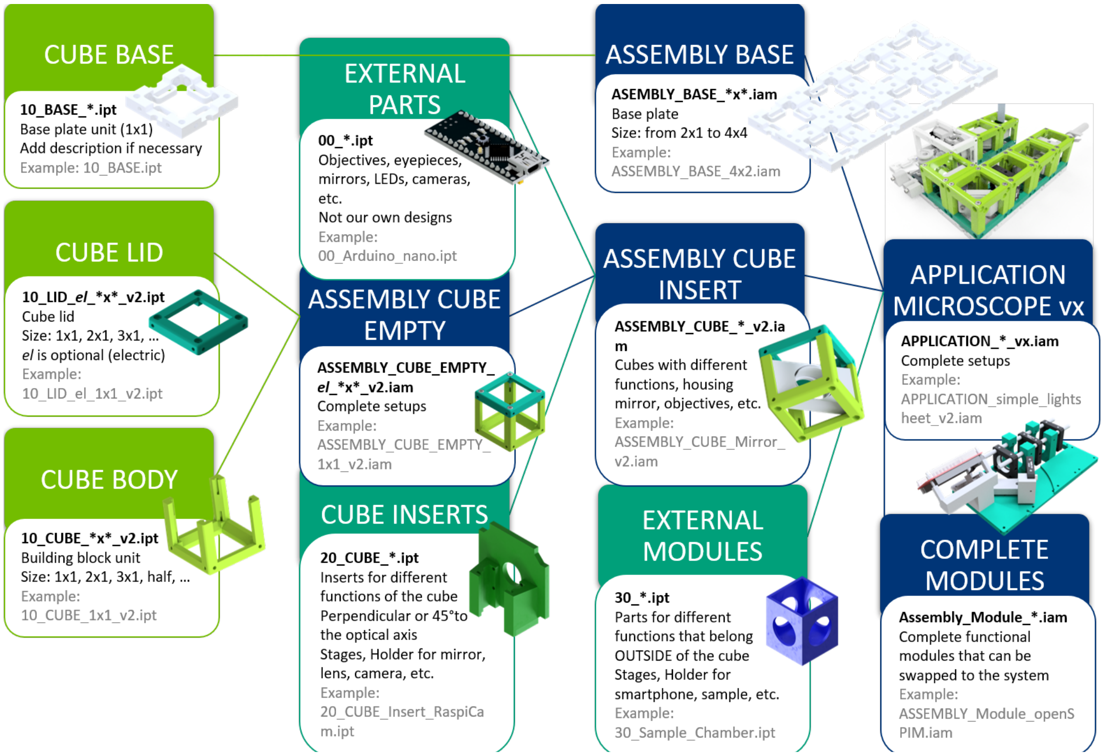

**Fig 4. The chart showing logical structure of building a UC2 setup.**

| Name | Description |
| --- | --- |
| **CUBE (Base)** | The units of the base are join into the **ASSEMBLY BASE**-plate ("skeleton" of the setups). This is the frame and backplane of the UC2 project, determining the size and layout of the optical system. UC2 modules attach to the rectangular baseplate positions using ball-magnets and ferro-magnetic screws. Additionally it supports smart-cubes with electrical power. |
| **CUBE (Body)** | The **ASSEMBLY CUBE (empty)** consist of the body- and the lid-part which gets screwed together. Screws can be inserted on all sides in order to build setups in three dimensions. |
| **CUBE (Lid)** | The **l**id closes the cube when attached by screw to the body. It can carry electronics like microcontrollers (e.g. Arduino, ESP32). By adding wires to the screws closing the **l**id electronic components can be supplied with electrical power. |
| **ASSEMBLY CUBE (empty)** | The raw cube/basic building block is made of the Body and the Lid and can vary in size (e.g. 1×1, 2×1, etc.). Note that the MDK only details the specification of the Cube and Base to the extent that it is necessary for module developers to develop modules. |
| **CUBE INSERTS** | Cube Inserts are physical components that implement various functions into the system by adapting EXTERNAL PARTS to the **ASSEMBLY CUBE INSERT**. They fit the inner dimensions of the cube. There are two types of inserts: perpendicular (to the optical axis) and diagonal. They serve as holder for various components like lenses, mirrors, cameras, filters, and other components demanded by the application. Existing inserts can be adjusted to fit specific parts (i.e. lens diameters). |
| **EXTERNAL PARTS** | Everything which is not part of the UC2-system or can not be 3D printed is termed as an EXTERNAL PARTS. This can be commercially available parts like objectives, lenses, LEDs, etc., but also 3D-printed parts from other projects (e.g. openflexure stage). |
| **ASSEMBLY CUBE (insert)** | This is the combination of the ASSEMBLY CUBE (empty) and a CUBE INSERTS. Since the ASSEMBLY CUBEs are the building blocks of a UC2 setup, adding features is accomplished by hardware plugins also called CUBE INSERTS. |
| **EXTERNAL MODULES** | Using EXTERNAL MODULES one adapt EXTERNAL PARTS that typically do not fit inside a cube but give function to it. This can be for example cellphones, stages projectors, etc. By providing customized hardware-adapter they interface with the ASSEMBLY CUBE. |
| **COMPLETE MODULES** | Entire Functional modules that can be swapped to the system. They have the correct screws and dimensions to adapt to the magnets on the baseplate. They are fully independent, but follow the optical path (e.g. SIM-module, ISM-module, projector, etc.). |
| **APPLICATION** | APPLICATIONs are complete optical setups or microscopes. They are composed of one or more base-plates (ASSEMBLY BASE) and modules with different functions. The github-repository provides a list of basic optical systems which are also compiled into a ready-to-use list called "TheBOX". |

#### The Cube

The cube is the cornerstone of the UC2 framework. Its purpose is to create a bridge between the toolbox and any external component which fits inside. It has a unit-size of $50\,mm$, with a hole-to-hole distance of $40\,mm$ and can be extended on an integer (i.e. $1 \times 1$ Fig. 5, $3 \times 2$, etc.) grid in all directions. The actual size is $49,8\,mm$ to incorporate imprecision of the printer. The center-symmetric cube is designed to provide guide the beam perpendicular and through the centre of the cubes' faces. The inner volume is large enough to adapt to common optical lab-ware (e.g. 1" Cage system from Thorlabs, Edmund optics, Qioptics etc.) and other components using customized adapters. Ferromagnetic worm and flat-head screws (DIN ISO 912, M3×18mm, DIN ISO906, M3×5mm, galvanized) also used to hold the cube together, connect to magnetically $5\,mm$ ball magnets sitting in the baseplate.

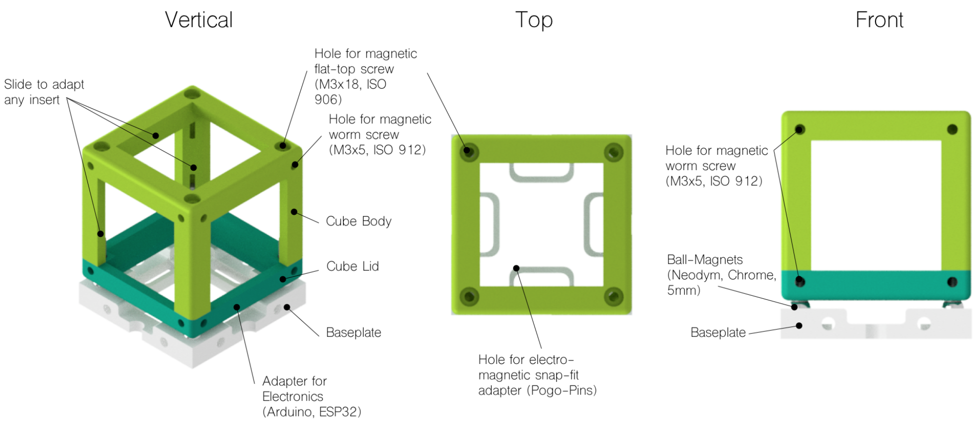

**Fig 5. Basic empty cube 1×1** The basic cube consist of two parts, the frame and the lid which is hold together by a set of ferro-magnetic M3 screws. These screws attach to the ball-magnets inside the base-plate. The inner structure of the cube allows inserting a customized hardware plugin in all directions.

#### The Baseplate

The Baseplate (unit-size $50 \times 50\,mm$, magnet-to-magnet distance $40 \times 40\,mm$, Fig. 6) is the "skeleton" of the UC2 framework and holds the different modules in place and provides a straight optical axis. The Neodym ball-magnets are press-fit into the 3D printed baseplate thus creating a stable mechanical connection to the 3D printed cubes. Knowing that it's mechanically over-defined, the 4-point interface is a compromise between a simple design-process for optical setups - which are aligned under 90° very often -, mechanical stability and versatility since cubes are much easier to stack than triangular pyramid or hexagonal units. Mechanical imprecisions e.g. due to faulty 3D printing can be compensated by adjusting the positions of the screws.
To provide electric components with power, wires added to the screws sitting in the cubes and to the conducting chromium ball-magnets can ensure an electric connection at a minimum number of visible cables since they are hidden inside the cube. In order to extend the grid in all room-directions the base-plate has holes at all faces to screw plates together. Additional M6 holes enables adaption to optical tables or breadboards to assure stable and long-lasting mount.

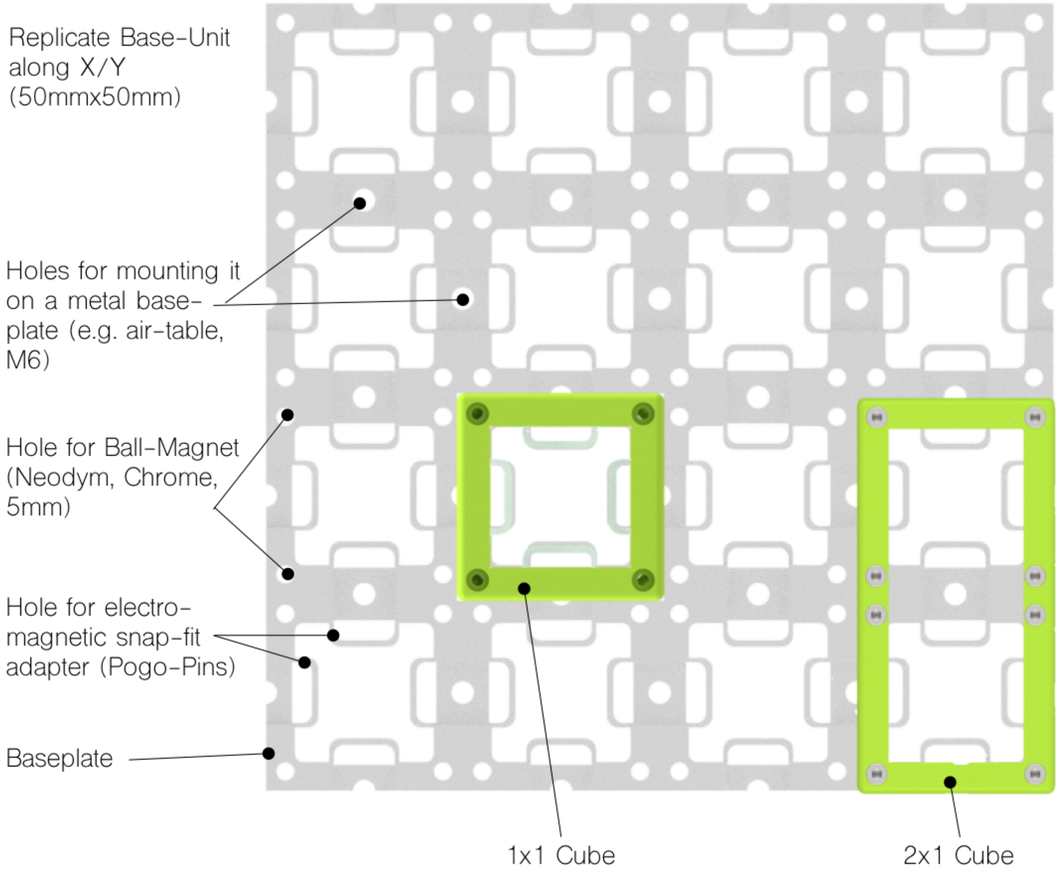

**Fig 6. Baseplate** $4 \times 4$ A exemplary assembly of a $4 \times 4$ baseplate equipped with two cubes. The M6 holes adapt to common optical tables to ensure long-lasting setups.

#### Cube Inserts

The cube inserts can be fully customized to adapt external elements, thus underlining the idea of creating an open-standard . The online-repository provides all relevant dimensions and CAD-Design templates for Autodesk Inventor and OpenSCAD to quick-start development with UC2. Additionally, a number of video-tutorials can be found in online video platforms. With this we invite people to develop their own modules which can contribute to the system.
Inserts are slided into the cube which allows variation in the position along the optical axis. By having dedicated rulers and spacers, one can make sure, that the insert is parallel to the cubes' face and optical dimensions can be reproduced. The two CAD-files below show examples for inserts at an angle of 0° and 45° w.r.t. the optical axis which fits into the standard point-symmetric $1 \times 1$-cube.

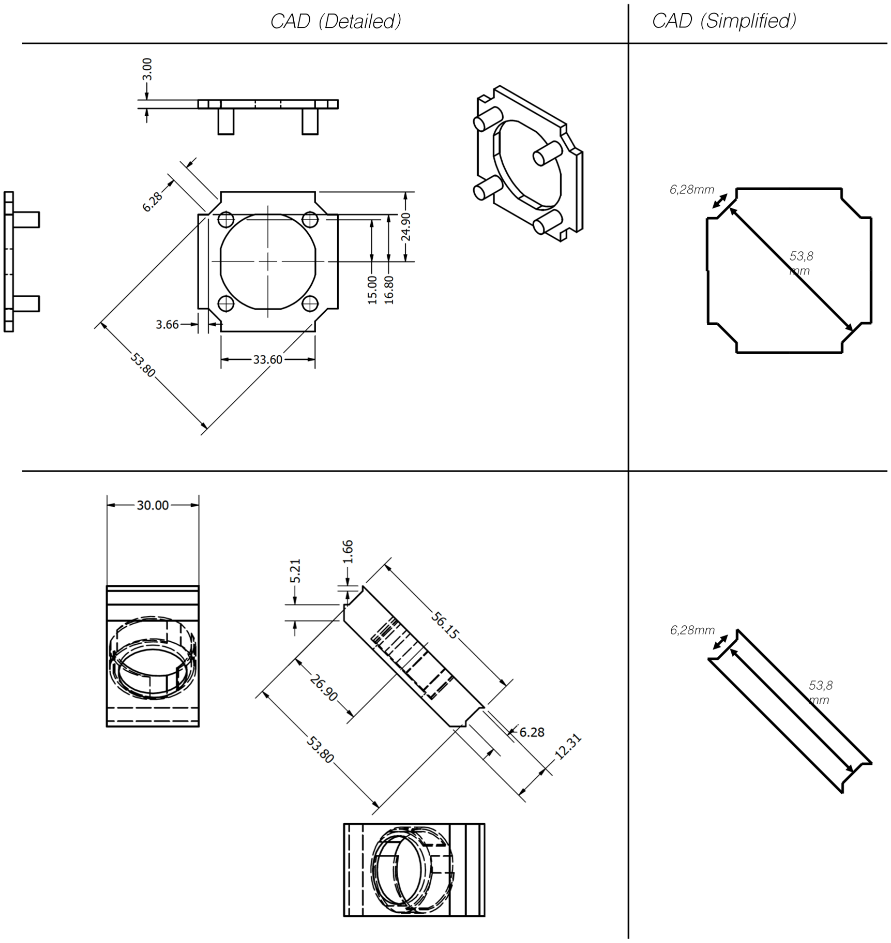

**Fig 7. Generic design for a cube-insert** - Since the cube is centro-symmetric, an insert can be rotated in all directions. The figures show exemplary insert-designs for a 0°- and 45°-version, meant for a thorlabs cage module and a mirror respectively. The smooth 3D-printed surfaces allow an easy sliding mechanism, but keep components in a fixed position in the same time. Newly designed cubes-adapter or inserts simply need to follow the dimensions are visualized in the simplified version of the CAD drawing.

#### Good Practice to transfer an optical system to UC2

The core idea of the modularity inside the UC2 system is based around the Fourier-optical principle, meaning that adjacent lenses are placed so that focal planes are interlocking in order to minimize effects like aberration and vignetting. This requires lenses to be an integer value of $50\,mm$ of their back-focal lengths, thus resulting in a focal-to-focal distance of about $100\,mm$. Determining Fourier- and image planes as optical interfaces enables sub-grouping of the whole system into modules and functional blocks. The optical axis always goes through the centre of and perpendicular to an open cube facet. Beam-folding in all directions (i.e. $X$, $Y$, $Z$) can be assured using mirrors.
In case of more complicated assemblies like the *open*SIM module, it is advisable to design a a monolithically printed block to assure higher precision and robustness. The outgoing plane (i.e. image for Fourier plane) should again adapt to the following plane from the next cube/module.

A simple example is given by a Keppler telescope illustrated in Fig. 8 which can be accomplished by concatenating two lenses ($f_1' = 50\,mm$, $f_2' = 100\,mm$) with a distance of $d_{1,2} = 150\,mm$ between their principle planes. The cellphone microscope shown in 8 gives another example how simple it is to create an imaging system, where the tube-length of typical finite corrected objective lenses (e.g. $d_{tl} = 160\,mm$) is reproduced by the two folding mirrors and a spacer before the intermediate image gets relayed by the ocular and imaged by the cellphone camera.

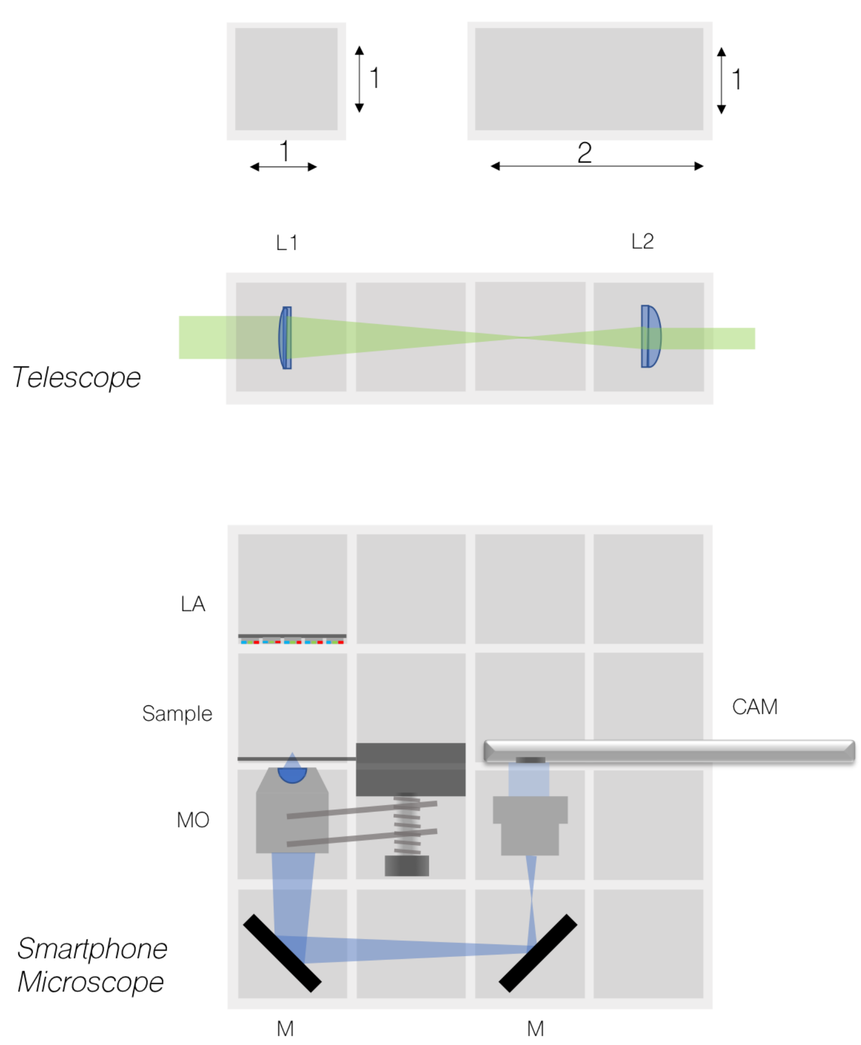

**Fig 8. Best practice for UC2 assemblies -** The core-units are the UC2 building blocks on a grid of integer $50\,mm$ (top). By combining two lenses $L_1, f_1' = 100\,mm$ and $L_2, f_2' = 50\,mm$ one can create a Keplerian telescope (middle); A more complex assembly can be created using objective lenses, LED matrices and oculars to create a smartphone microscope (below).

#### Software

#### Details about Hardware and Software Control

To create reproducible long-term measurements with the incubator microscope, we created Python-based GUI which runs on a Raspberry Pi equipped with a 7-inch touch-screen. A detailed description on how the system need to be installed can be found in the dedicated Software-Repository. The user interface written in Python and based on the kivy-framework [1] is visualized in Fig. 9 and allows the control of several hardware elements such as individual addressing of LEDs in the LED-matrix, movement of motors connected to the system (e.g. X, Y, Z) and intensity control of the fluorescent illumination. In addition to that, the software also allows the scheduling of long-time experiments. This includes the choice of the illumination modality (e.g. DPC, Fluorescence, bright-field, dark-field,etc.), the periodicity of frame-acquisition and the whole duration of the experiment. The images captured using automatic settings such as auto-exposure and auto white balance (AWB), are saved as JPEG-compressed photos on the internal SD-card in order to save memory.

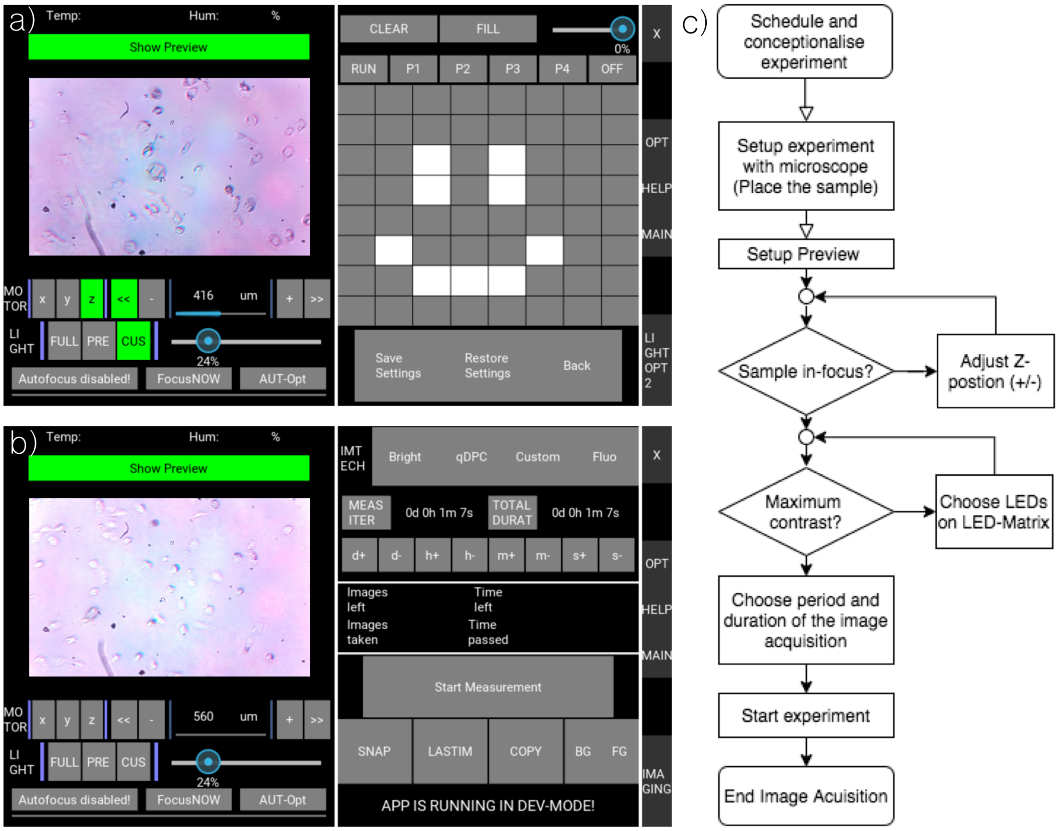

**Fig 9. Basic settings for the GUI** The GUI is divided in the hardware-control section* a) and experiment configuration panel which to setup long-term time-lapse series e.g. for the incubator or light sheet microscope. In c. an exemplary workflow of a typical biological experiment over multiple days is visualized.

The software can be used to control wired as well as wireless components connected to the Raspberry either by $I^2C$ or WiFi. A dedicated UC2-$I^2C$-device adapter created in Python as shown in Fig. 10 preserves the modular nature of the UC2 system since the command-set sent by the Raspberry Pi to any $I^2C$ or MQTT device in the same network follows a modular type-set.

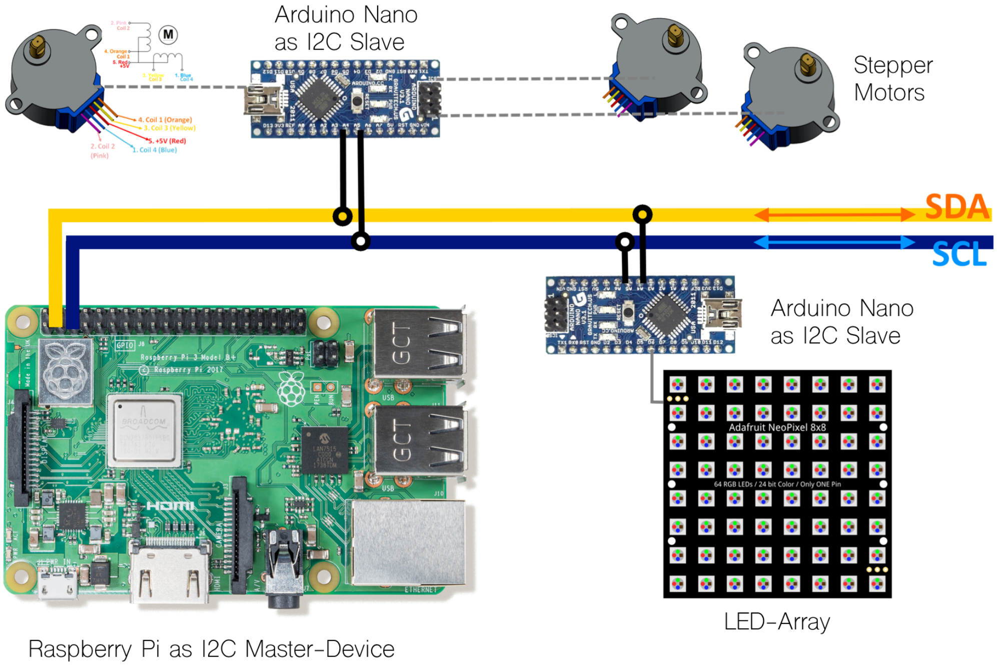

**Fig 10. Schematic of the $I^2C$ device adapter** The Raspberry Pi acts as a $I^2C$ master device which sends controlling commands to all slave in the same network created by the four-wired signal (*5V:* power-signal, *GND:* ground-signal, *SDA:* signal data, *SCL:* signal clock ). UC2 relies on low-cost Arduino Nanos which convert the $I^2C$ commands into hardware control operations for motors, leds or anything else controllable through microcontrollers.

The MQTT-based wireless system visualized in Fig. 11 has the advantage, that multiple devices can overtake the control of an optical system, beneficial for example in cases one cellphone acts as the image-acquisition device and another cellphone is used as a remote-control for a setup. Since the devices can be reached from remote places through the internet, adjusting or readout of parameters could theoretically be done from any place which supports internet access. This feature has not yet been tested in order to follow the internal privacy policy.

Best practice for the TCP-IP based MQTT-network connection is to setup a dedicated WiFi-Router (Netgear Nighthawk R7000) which handles the different connections. A MQTT broker (i.e. server) can be created using either a cellphone or the Raspberry Pi by using open-source software such as Moquette or Mosquitto. We also developed a stand-alone Android APP which incorporates the MQTT broker as well as the MQTT client in order to use the system independent from any external devices (e.g. in the fied). The source-code can also be found in our Software Repository

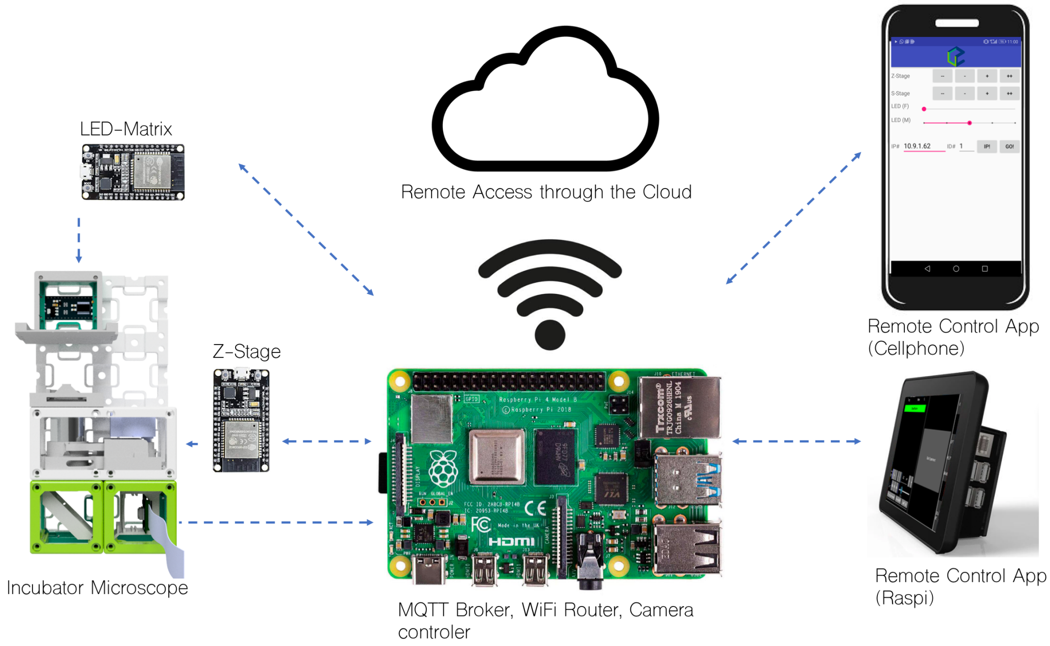

**Fig 11. Schematic of the MQTT Connection** All devices are connected to the same Network (e.g. WiFi hotspot) and MQTT-broker (e.g. server) which can be represented by a Raspberry Pi. The MQTT-based network protocol allows multiple devices to be used remote controls. Each MQTT client (e.g. ESP32) reacts on a sent command.

#### Experimental Details

#### Long-Term In-Incubator Microscopy

The aim of this biological study was the long-term observation of macrophages in vitro under different environmental effects. Other than putting an incubator on a microscope stage, we decided to put the whole microscope in a bench-top incubator (Heraeus Instruments, Germany) which ensures suitable conditions for living organisms (e.g. Temperature, $CO^2$-level). We formulate requirements for long-term biological imaging as follows:

- Time-lapse imaging at one-frame per minute over several days
- Optical resolution on cellular level (e.g. $2 - 3\mu m$)
- Autofocus to compensate drift of the sample due to temperature-depending deformation of the plastic microscope along $z$
- Bright-field and fluorescent imaging of labelled cells (e.g. cell-tracker green)
- Customized illumination settings to enhance the visible contrast of weakly scattering phase objects
- Autonomous operation over long-time periods
- Host standard microscope slides and 35 $mm$ Petri-dishes

The resulting prototype which was created based on these requirements using the UC2-system is shown in Fig. 12 and in more detail in the online repository. It follows a simple optical path derived from an inverted compound microscope with a finite-corrected objective lens (Generic brand, 10×, NA=0.3) following in a theoretical resolution of $1.8\mu m$ with an coherently

illuminated sample (e.g. only one LED). In order to reduce the overall size of this device, we reduced the tube-length from $160\,mm$ to $\approx 100\,mm$ which follows in a shorter working distance and reduced effective magnification. The resulting optical resolution is $d_{min} < 2.3\,\mu m$ quantified with imaging a USAF chart visualized in Fig. 13 resulting in an effective magnification of $\approx 7\times$ on the Raspberry Pi camera (V2, Sony IMX 219, $d_{pixel} = 1.4\,\mu m$, Bayer-pattern, $t_{exp} = 100ms$, UK, v2). To achieve multi-modal imaging, we used a $8 \times 8$ RGB LED array (Adafruit #1487), where only a subset of the available LEDs are within the NA of the detecting objective lens. The selection of the LEDs was done through a customized GUI on the Raspberry Pi while the visible contrast was maximized. For fluorescent illumination we decided to use a dark-field like illumination from below the sample. A module, sandwiched between the objective lens (e.g. Z-stage) and the sample, hosts a number of high-power LEDs sitting on a star-LED, while the resulting dark-field illumination blocks the zeroth-order which makes selecting the emission filter more cost-efficient, since only the residing thus weaker stray-light has to be filtered out.
All electric components are connected to a micro-controller which was an Arduino Nano in the wired (e.g. $I^2C$) and an ESP32 in the wireless (e.g. MQTT) control-mode. The magnetically hold LED-matrix can be dismounted to have more space during media change or in case malfunctioned hardware needs to be replaced. Being free in the choice of the distance between the sample and the illumination unit gives additional space for wires and tubes for applications like flow-cytometry or lab-on-the-chip.

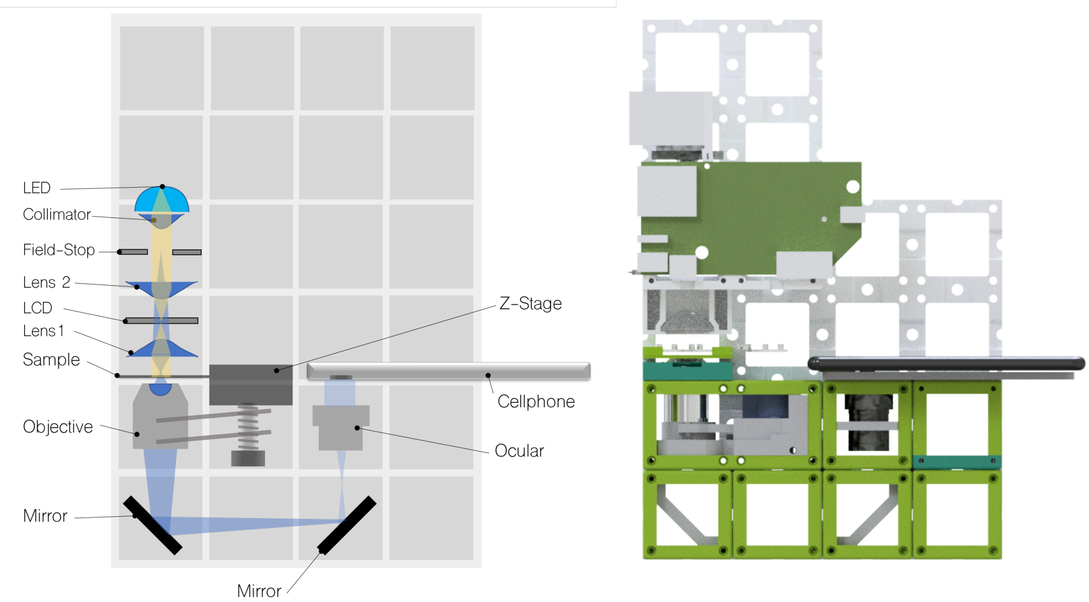

**Fig 12. Scheme of two inverted microscope used in the incubator** - A LED-array allows the selection of the illumination angle and enables quantitative imaging. The objective lens inside the Z-stage can be moved up- and down by a stepper-motor controlled by an ESP32, while the optical path is folded using a simple mirror to form an image on the Raspberry Pi camera. The camera is connected to a Raspberry Pi equipped with a 7inch touch-screen which gives an overall price-tag of $\approx 300$ Euro

To be able to focus the sample during the acquisition series we designed a customized monolithic Z-stage (see our github-repository) which is inspired by the open-flexure design Bowman et al. [15]. It is based on a spiral-bearing where a level-arm pushes the objective up and down using a stepper-motor (China, 28BYJ-48). This way, the Z-stage observes no radial shift while it's moving. To make sure, that the sample stays in place, it is fixed using a magnetic clamp which makes removing the sample in case of exchange of the cultivation media easier. An additional module which hosts a pair of low-cost stages allows movement of the sample in XY (see xyz-assembly)with a precision of $\approx 20\,\mu m$. To keep the experimental setup simple, we forgo this feature.

In-vitro measurements where achieve by putting the microscope into a standard S2 biological laboratory, while all parts where sprayed with 70 % ethanol before entering the lab area. After setting up the microscope, the imaging parameters are selected and the microscope runs for several days before the data gets transferred from the Raspberry Pi to an external storage media for further processing.

For the details about cell-preparation, please have a look in the section* 3.

The $8 \times 8$ RGB LED array also allows quantitative imaging based on the work by Tian&Waller[17] which captures a series of obliquely illuminated phase-objects and performs a deconvolution with the corresponding transfer-functions. This feature was not used during the long-term acquisition since the contrast was already satisfactory and the additional effort to compute each result-image was rather high. To track changes in the the long-term in-vitro experiments with low-contrast cells (e.g. unlabelled macrophages) we relied on oblique illumination to promote the phase gradient of the mostly phase objects.

All design-files including the bill-of-material ($\approx$ 300$)and an illustrated step-by-step assembly tutorial can be found here.

**Optical Resolution and long-term stability** Since the very first experiment in the biological lab, the cube-based design went through a a series of iterative optimizations, where certain components were exchanged and optimized over time. The portable design of the microscope simplified the transportation (e.g. using a bike, see Fig. 14) from the workshop to the University Clinic Jena (UKJ), where the experiments were performed. Shaking and heavy vibration did not alter the quality and stability of the setups underlying the use in the field. During the development cycles of the BF-microscope we had to change system-paradigms multiple times and hence brought the systems back from the university hospital to our optical labs and vice-versa, meaning roughly 6km per direction. We directly used the transportation as a stress-test of system-robustness by only roughly packing it into a bag and then carrying the light-weight systems by bicycle. Even after many transports, the systems are still taking images of identical quality.

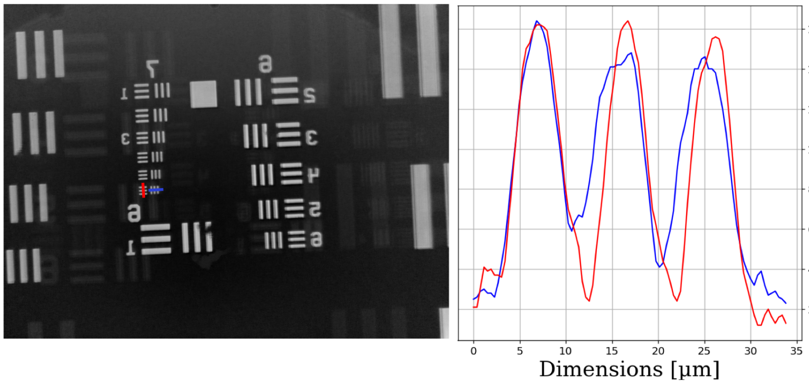

**Fig 13. Calibration using the USAF** - The incubator microscope can resolve sub-cellular features demonstrated by imaging Group 7/Element 6 of a USAF 1951 chart (Thoralbs, R3L1S4P, USA) resulting in a resolution $d_{min} < 2.19\,\mu m$.

Nevertheless, we want to emphasize, that due to the use of 3D printed thermoplastic material (PLA, ABS), certain parts tend to bend during long-term experiments. By choosing ABS over

PLA everywhere where components experience larger tension, like the z-stage and the base-plate, the problem can successfully be compensated. We found, that once the Z-stage settled, it experience almost no deformation over long-time periods which makes adjustments unnecessary. One stage equipped with a 10×, NA=0.3 objective lens was in-focus even after 3 month.

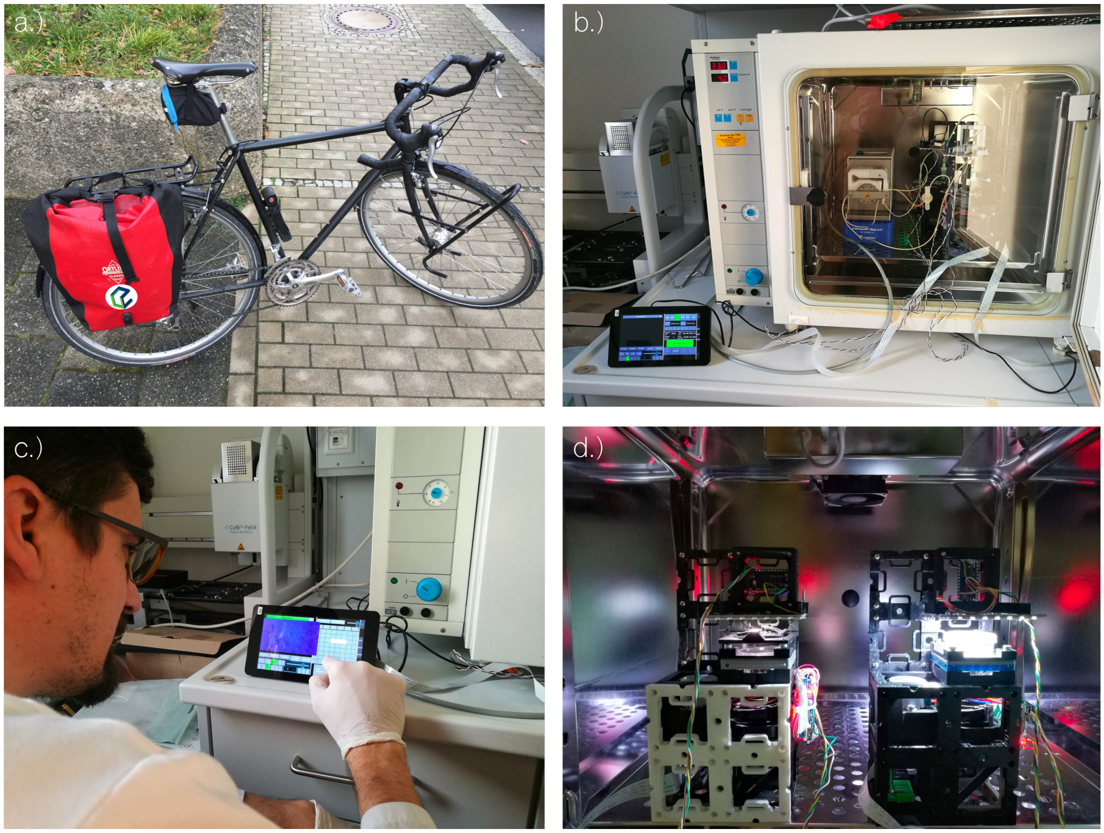

**Fig 14. Setting up the incubator microscope -** a) The microscope fits inside a small box and can conveniently be printed and assembled at one point and transported to the bio-lab using a commuter bag on a bike. In b) we show a customized application where a Microfluidic Ibidi µ-chip with endothelial/macrophage perfused co-culture is placed on the incubator microscope, before all cables are connected. c) The next step requires setting up experimental details such as duration and interval, as well as illumination settings can be set using the touchscreen on the GUI. d) due to their small footprint multiple dives find place in one incubator and can conduct multiple experiments at the same time.

Especially in long-term experiments it is of great importance, that vibrations are minimized. The here used bench-top incubator (Heraeus Instruments, Germany) was placed on an ordinary lab-bench which experiences low-frequency vibrations resulting from footsteps. The resulting fluctuations of the FOV in the long-term experiments (see Supplement 2) can partially be compensated using a heavy metal-plate as a base for the microscope during the experiments or using a image registration routine in the image post-processing.

#### Quantitative Phase: aIDT

To give an example that our modular optical system can be used with a variety of different open-source image processing algorithms, we choose the freely available code for the annular intensity diffraction tomography *aIDT* from Li et al. [9]). The algorithm is especially interesting since it only requires a series of images with a varying illumination direction $k_{illu}$, where the algorithm can self-calibrate the illumination direction - ideal for a system which may experience slight misalignment over time. Other than methods like Fourier Ptychography Microscopy (FPM), the detection requires only illumination angles close to the edge of the detection pupil (e.g. dark-field illumination) . Therefore we add an RGB LED-ring (Adafruit, #1643), where each LED can be addressed individually using a microcontroller (e.g. Arduino Nano, Espressif ESP32). We used only the green-channel to produce quasi-monochromatic light and acquire a set of images of fixed endothelial cells using a cellphone (Huawei P20, BI-CMOS Sony, IMX 286, China). It was of great importance to acquire the data in RAW-mode since the automatic calibration routine of the *aIDT*-algorithm fails when the images are compressed (e.g. JPEG).

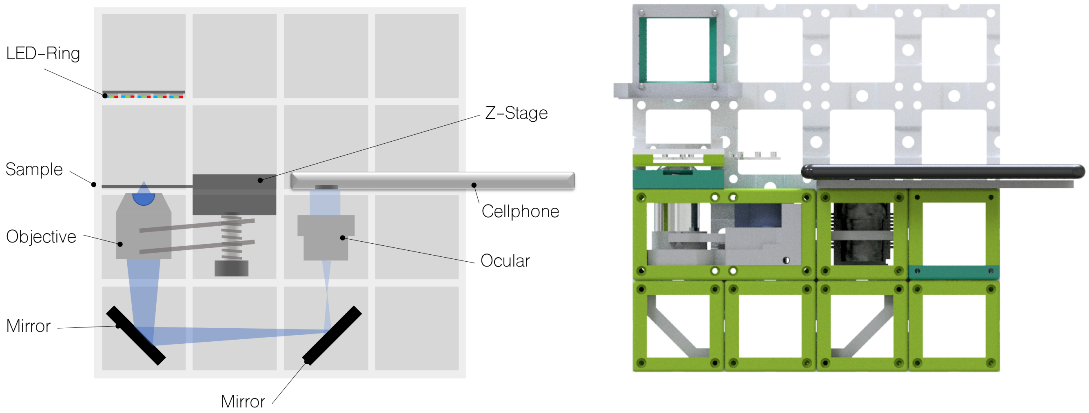

**Fig 15. Scheme of the aIDT assembly using a LED-ring and cellphone camera -** The LED-ring illuminates the phase-sample from 16 different angles which produces a series of images feeding the inverse model. This algorithm can recover a focus-stack of the quantitative phase. The cellphone can send MQTT commands to the LED-ring to synchronize illumination ⇄ frame-acquisition.

The optical design follows the one of the In-Incubator microscope described in section* 3, where we used a 10×, NA=0.3 objective lens. The illumination NA has to be slightly less than the detection NA to have all illuminating plane wave inside the pupil which follows into the requirement of $NA_{illu} \leq NA_{det}$. Since the LED-ring has a radius of $r_{ring} = 16, mm$, the $NA_{illu}$ is governed by the distance between the LED and the sample $d_{sample}$:

$$NA_{illu} = tan(r_{ring}/d_{sample}) \quad (1)$$
$$d_{sample} = r_{ring}/atan(NA_{illu}) \quad (2)$$

which follows in a distance $d_{sample} \geq 54\,mm$ and adjusted experimentally to about $74\,mm$ which follows in a smaller effective NA. Therefore, an additional layer in the base-plate (not shown in Fig. 15) was added. A modified version of the original code along with all necessary design files and manuals for this experiments are published in the github-repository.

#### Selective plane illumination using a Light sheet microscope

Even though the term "Ultramicroscopy" [16] has been around for almost a century, the topic selective plane illumination microscope (SPIM) or light sheet microscopy (LSM), where a thin light plane illuminates a (fluorescently labelled) sample perpendicularly to the detection direction, gained lots of attention during the last decade. It provides gentle 3D-imaging of volumetric in-vivo samples [13, 14]. Though this concept of optical sectioning in order to increase the optical resolution along the detection axis is straightforward, it becomes even more obvious if one experiences it in a hands-on experiment. Therefore, we started a series of workshops to demonstrate the working principle of these microscopes, available with a comprehensive alignment tutorial in online repository. The overall *open*SPIM-inspired setup visualized in Fig. 16 is kept simple in order to give users the chance to build these setups on their own.

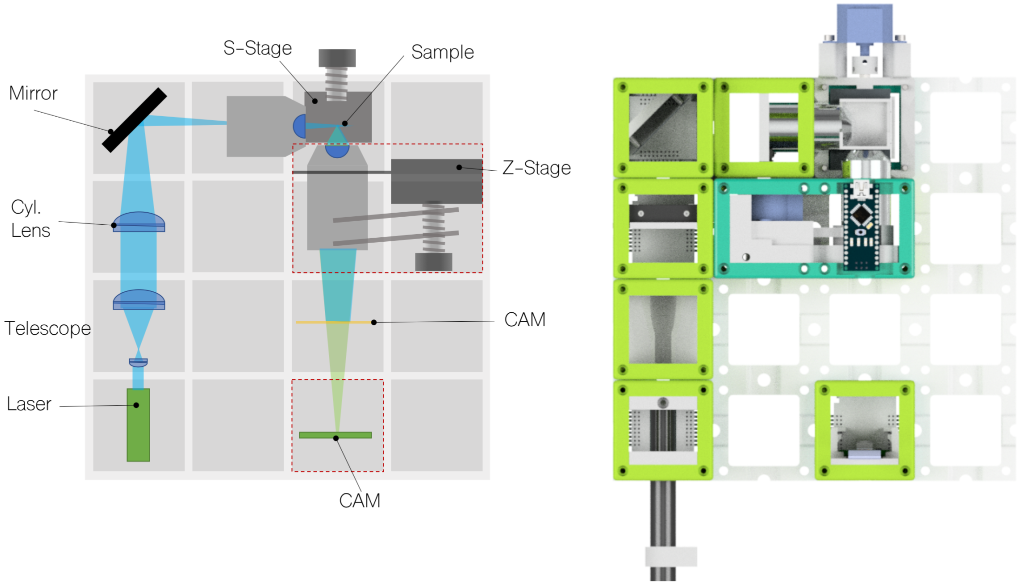

**Fig 16. Scheme of the selective plane illumination setup** - The left schematic shows the optical diagram of the SPIM, where a laser-pointer is first expanded and then focussed by a cylindrical lens into the BFP of the illuminating objective lens. The detection is accomplished with an finite-corrected microscope perpendicular to the illumination plane (e.g. light sheet). The sample is placed on a z-stage which can perform a focus stack of samples placed in a water chamber. A kinematic mirror mount can be used to align the light sheet.

**Optical setup** The setup hosts a blue laser pointer ($\lambda_c = 445\,nm$) as the illumination source which gets expanded by a telescope. This telescope first focusses the incoming parallel light using a cellphone lens (Apple, iPhone 5, NA=.24, $f' = 3.2\,mm$, $\approx 5\$$) before it gets collimated by a second lens ($f' = 20\,mm$) to achieve a magnification of $\approx 6$. This beam gets partially focussed by a cylindrical lens (Comar optics, $f' = 63\,mm$) to create the 1D line-profile before it passes a kinematic mirror mount cube accomplished by ball-magnets sitting on 3 ferromagnetic M3 screws, followed by a magnetic plate (e.g. galvanized steel, $30 \times 40\,mm$). This way the beam can be precisely placed in the BFP of the illuminating objective lens (e.g. 4×, NA=0.14). The resulting light sheet inside the sample plane has a theoretical thickness of 200 $\mu m$ based on rather pessimistic assumptions on the profile of the laser-diode, while the actually measured thickness is around $45\,\mu m$ and thus slightly better than the depth-of-field (DOF) of the objective lens $d_z \approx 56\,mum$. A clear advantage in this configuration is given by the ability to scan the light sheet in order to generate a volumetric image stack. This can be achieved by either moving the kinematic mirror or moving the sample. When changing the angle of the mirror, the light sheet can scan along the optical axis of the detection objective lens. Since the DOF is rather large, most of the object information are in-focus, while the light sheet excites only those fluorescent labels within its beam volume. When using a static light sheet, aligned onto the in-focus plane of the detection path, the z-stack is obtained by moving the stage that carries the sample and acquiring an image for each step.

The sheet illuminates the fluorescent sample sitting on a movable sample-stage. The sample-

stage is equipped with a stepper motor (China, 28BYJ-48), which pushes a magnetic plate sitting on a flexure bearing. The step size is governed by the pitch of the screw and the smallest step size of the motor which leads to a reproducible step size of $\approx 10\,\mu m$.The sample holder directly usable for syringes holding samples fixed in agarose, is equipped with 3 ball-magnets to move the sample coarsely to get a roughly focussed image. Additionally, a 3D-printed water chamber can be placed on the moving sample stage to reduce scattering and aberration of the illuminating as well as the detection beam path.
The detection path (Fig. 16, green) follows a typical compound microscope as also illustrated in Supplementary 3, where either a 4×, NA=0.14 or a 10×, NA=0.3 objective lens were used. As the detector we choose either be a Raspberry Pi camera module without a lens or a Raspberry Pi/cellphone camera with a lens but combined with an eyepiece (20×). As an excitation filter we relied on a gel-filter (Lee, #010, medium yellow).
Acquiring a z-stack can conveniently done using the GUI running on the Raspberry Pi. It automatically moves the sample by one step and acquires an image for $N$ z-positions.

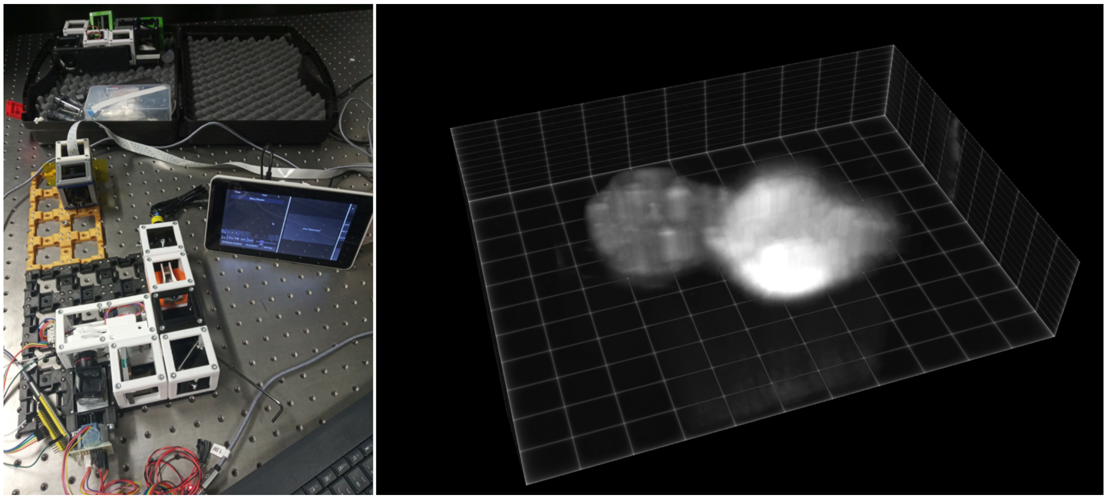

**Fig 17. Light sheet microscope** Left - the complete setup. Right - 3D reconstruction of a zebra fish embryo head from a z-stack obtained with this setup.

**Alignment of the setup** We provide a detailed description of the alignment procedure in our github-repository. Additionally the Video Supplement 2 gives an introduction on how to convert the incubator into a light sheet microscope within 5 minutes.

#### Image Scanning Microscope (*open*ISM)

Many consumer-grad electronics like video-projectors (e.g. digital mirror devices, DMD) or movie screens enabled entertainment "on-the-go" for a very low price - compared to scientific instruments - due to mass-production. Besides wide-field projection systems based on liquid crystals on silica (LCoS), LCD or DMD displays, more exotic laser-scanning based systems (e.g. Sony MP.CL1A, Japan) enabled us to create a UC2-ready image-scanning microscopes (ISM) for around 300 Euro.
The laser scanner, equipped with a small Micro-Electro-Mechanical System (MEMS) scans a set of RGB ($\lambda_{blue} = 450nm$, $\lambda_{green} = 530\,nm$ and $\lambda_{red} = 650\,nm$) laser-beams over the 2D (e.g. X/Y) plane with a frame-rate of $60\,fps$ at a spatial resolution of $1920 \times 720\,pixel^2$ to create an aerial image. A customized UC2 module enables the integration to our $50 \times 50\,mm^2$ standard. Following the work by Enderlein et al. (ISM) [11] and York et al. (iSIM)[18] we illuminate the sample with a nearly monochromatic, non-overlapping lateral grid of laser-light. The pattern

then gets translated along a unit vector in X/Y such that the sum of all illumination-patterns corresponds to a laser-scanning bright-field illuminated image. Each grid-point can be interpreted as a confocal-pinhole and thereby sectioning is improved as compared to standard bright-field imaging.
In Post-processing, all illuminatination spots (per frame) are treated in parallel. For each spot, a tile - meaning pinhole of multipixel size - is placed around its centre and gets extracted. Non-centre pixels then get moved to center by half its distance to account for the most-probable position of the fluorophore with respect to the difference between illumination and detection pixel. Finally, the signal is integrated and written into the final image at the position where the tile-centre was placed. This procedure leads to a resolution of a factor up to $\sqrt{2}$ [10] compared to standard confocal microscopy.

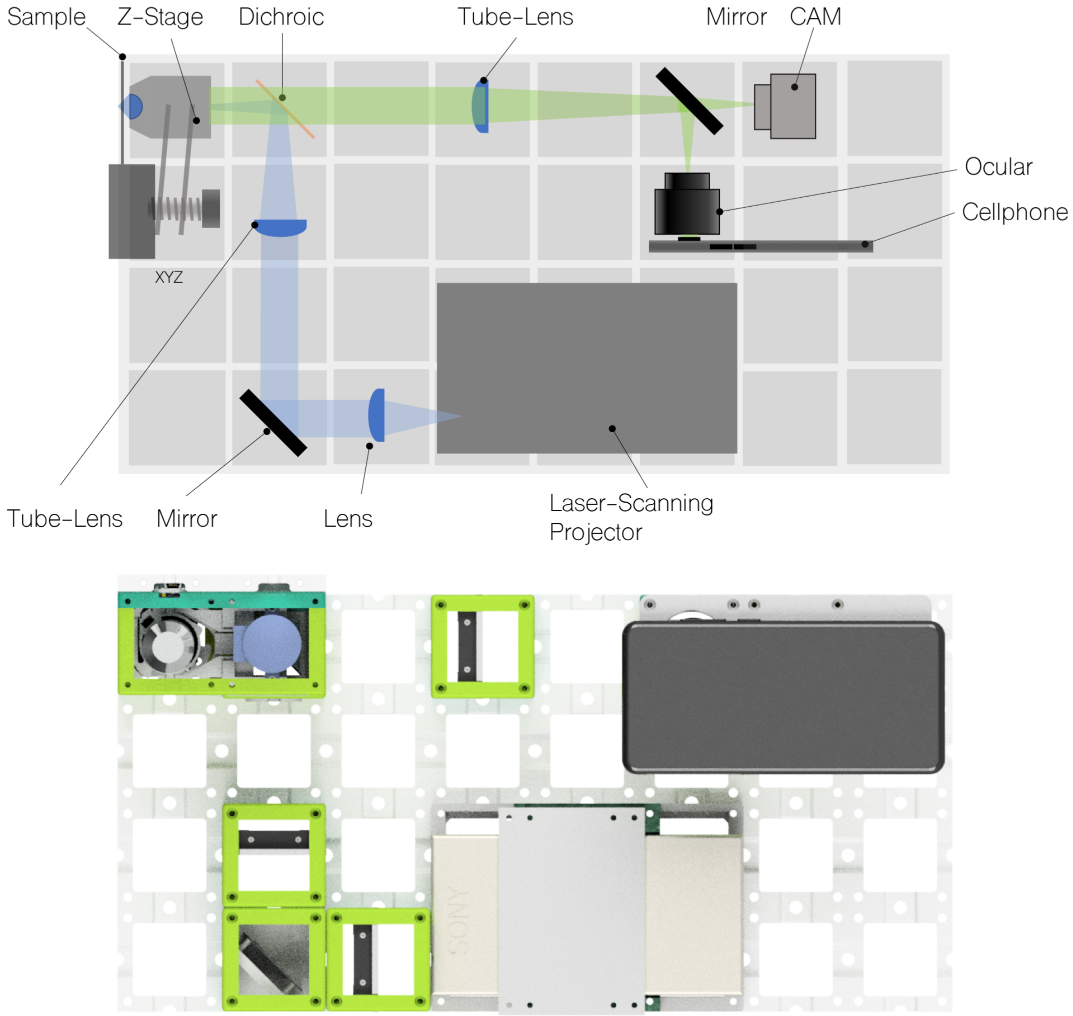

**Fig 18. Scheme of the image scanning microscope (openISM)** - The light-path shown in the schematic above starts with the laser-scanning projector, where the beam gets collimated using the lens $L_1$ and re-imaged into the BFP of the objective lens using Lens $L_2$. This telescope magnifies the mirror by a factor of 5. The detection path (green) follows a typical infinity-corrected microscope where either a CMOS (e.g. IDS, BASLER) or cellphone-camera combined with an eye-piece.

The design-files and additional explanation can be found here: here.

**Optical setup and frame acquisition** The optical setup shown in Fig. 18 is straight-forward. The resonating MEMS in the projector needs to be imaged into the BFP of the microscope objective lens. In order to get high-resolution images, the BFP is over-filled by the mirror ideally. We assumed a diameter of the aluminium mirror of $d_{mirror} = 1.5\,mm$ and a diameter of the BFP of around $d_{BFP} = 5.5\,mm$ which requires a telescope, created by a lens $f'_1 = 30\,mm$ and a following tube-lens $f'_2 = 180\,mm$. The low-cost infinity-corrected microscope objective (Optika, 20×, $NA = 0.4$, N-plan) was put in a motor-driven Z-stage to allow focussing the objective relative to the sample. A set of different dichroic-mirror cubes with suitable filters (excitation/dichroic/emission-filter: Comar Optics, 465 IK/510 IY/526 IB) allows the switching between different fluorophores and excitation wave-lengths. The detection path was generated by a $f'_{TL} = 180\,mm$ tube lens before a 20× mono ocular propagates the image to infinity. This way a cellphone-camera can create a sharp image if the autofocus is set to infinity. The effective-pixel size depends on the selection of the cellphone and results in $d_{pix} \approx 150\,nm$ when using the Huawei P20 Pro.
Since the laser-scanner is not meant to be used for scientific instrumentation, technical details are hardly available which makes interaction with it cumbersome. Also, the uncommon resolution of $1920 \times 720\,pixels^2$ follows in an unknown interpolation of the image sent by ordinary computers, thus not showing the "true" pixel information (i.e. one-to-one pixel relationship). We solved this by using a Macbook (Apple, 13-inch, MPXQ2D/A, USA) using an USB-C to HDMI adapter at a display-resolution of $720p$ in combination with a customized Python-script which generates and displays the ISM-patterns. The monochromatic cellphone-camera (Huawei P20 Pro, China) was driven using the open-source software FreeDCam ([4]), where the exposure time $t_{exp} = 1/60\,s$ matches the frame-rate of the laser-scanning projector in order to reduce temporal bouncing effects of between frame rate and laser round trip.

#### Structured Illumination Microscopy (*open*SIM)

A dedicated SIM-module, based on the recent publication from Sandmeyer et al. [**Sandmeyer2019**] integrates a low-cost single-mode diode laser ($532\,nm$) and a Raspberry Pi driven DMD module into the UC2 system in order to generate a sinusoidal pattern for super-resolution microscopy. The monolithically printed module (Fig. 19) which directly adapts to the magnetic grid-system, makes sure that all components are correctly aligned when mounted in their predefined positions (i.e. mechanical stop). This simplifies the alignment process dramatically. By addressing a set of gratings on the DMD (Fig. 19), relying on the coherent illumination of this pattern and Fourier-filtering the zeroth-order, a two-beam interference pattern is formed within the sample plane. Using proper image processing, a set of 9 images, where the grating's-phase and its orientation is varied, can potentially increase the lateral and axial resolution when fluorescent samples are used [**Wicker2014a**]. With this module we are not yet aiming for a gain in optical resolution, but rather for the use in educational areas and for rapid prototyping where data generation (e.g. machine-learning) or development of image processing algorithms is of interest. The design-files as well as the bill-of-material and detailed instructions on how to build this module can be found in the online repository. The cost of this module is in the range of 300 $.

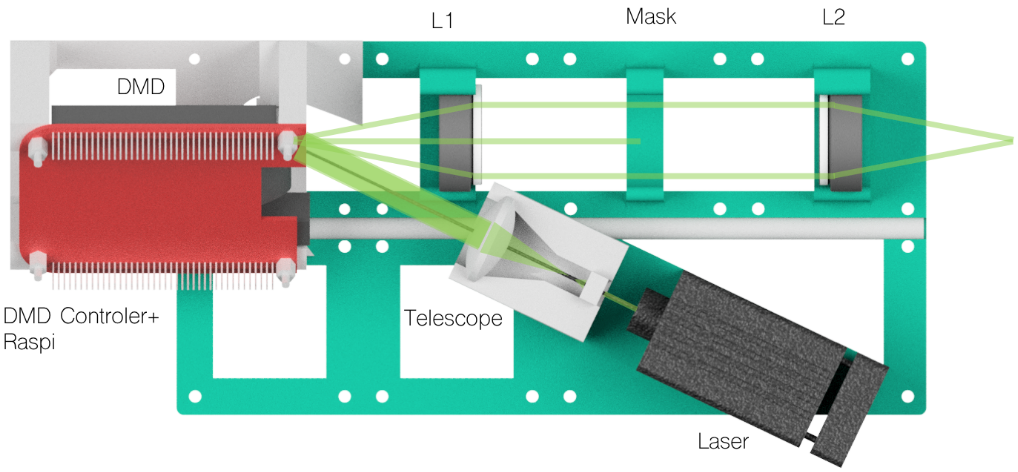

**Fig 19. Ready-to-print *o*penSIM module** - The components are directly mounted on a monolithically printed frame which holds the Laser, an adjustable beam-expander using a kinematic mount, the DMD + Electronics and the telescope which relays the digital grating and filters the Fourier-plane

The design-files and additional explanation can be found here.

**Optical setup** The optical setup of a classical two-beam interference SIM [7], also visualized in Fig. 19, is straight-forward and based on a set of $4f$ systems, thus ideal to reproduce with the UC2 system. The structured illumination (i.e. sinusoidal pattern) follows from a two-beam interference using two plane waves reaching the sample under an oblique angle. A super-resolution effect is achieved, when the cut-off frequency of the resulting grating is close to the highest possible frequency transfer of the used detection lens, which necessitates to place two delta-peaks which are capable to interfere (e.g. temporally/spatially coherent with each other) close to the rim of the circular BFP.
In order to have a more compact system which can be transported to different educational events, the DMD-imaging telescope is based on two lenses with $f' = 50\,mm$ (e.g. Thorlabs ACS-254 50). A tube-lens of $f' = 180\,mm$ images the spectrum of the DMD (Texas Instruments, DLP2000EVM) previously filtered by a customized and 3D-printed Fourier-mask to block the zero-order into the objective lens's BFP. A set of silver and dichroic mirrors fold the beam-path as indicated in Fig. 20. Flipping the DMD by 45° around the optical axis and introducing an angle of 25° between the DMD and the illuminating laser-beam, takes the blazed-grating effect of the micro-mirror device into account and therefore maximizes the grating contrast and intensity in the $\pm 1^{st}$ diffraction orders.
The DMD is driven by a Raspberry Pi using a customized PCB and displays the 9 patterns which have been generated using the open-source software suit fairSIM [12].
For the detection, one can choose between a standard industry-grad CMOS camera (e.g. Basler, IDS, etc.) which is placed in the intermediate image after the tube-lens or a cellphone-camera, where an additional eyepiece (Müller, 10×) was used to relay the intermediate image to the entrance pupil of the cellphone lens. Correct imaging in this case is assured if the Ramsden disk of the ocular (e.g. exit pupil), matches the entrance pupil of the cellphone.

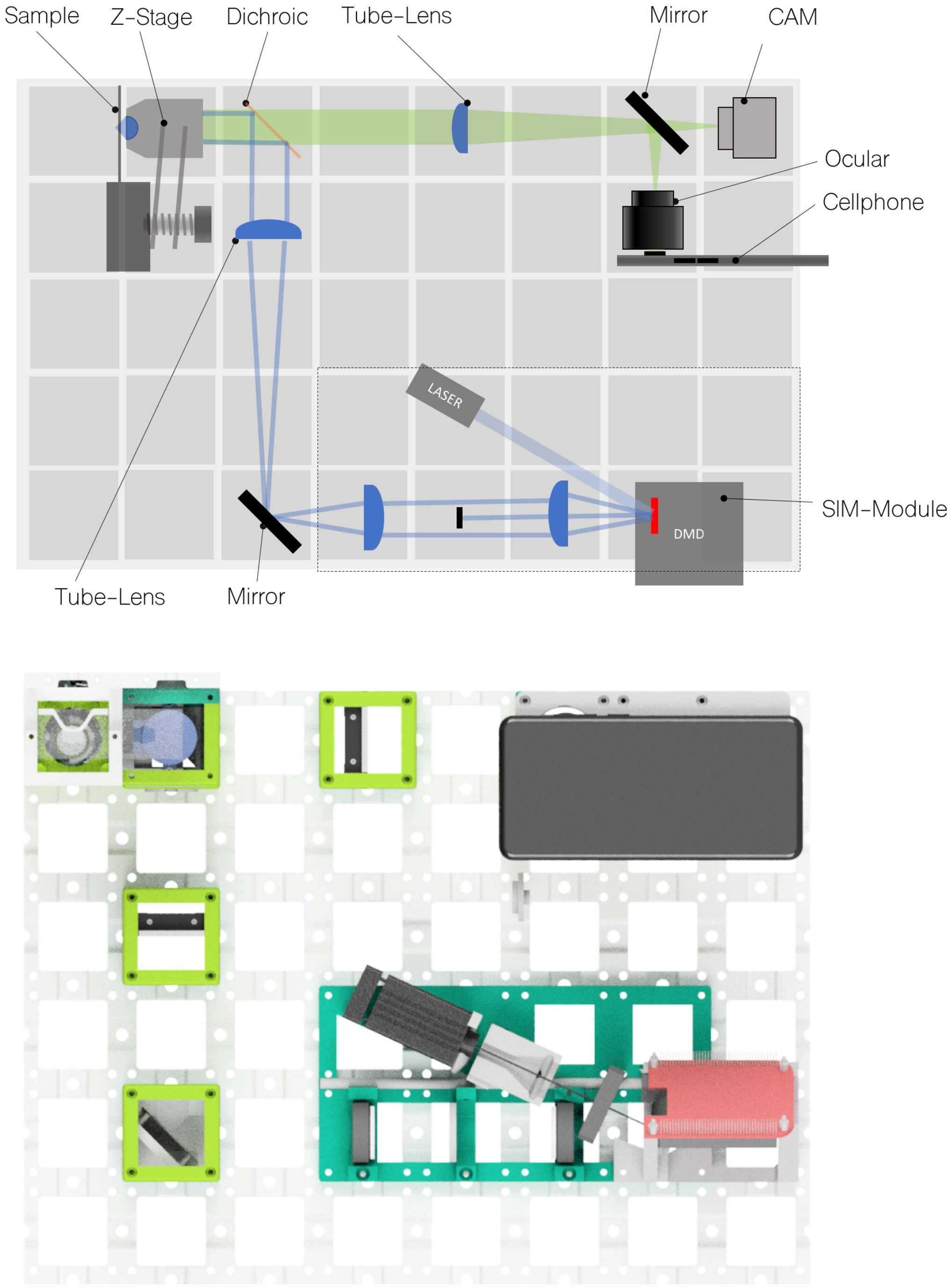

**Fig 20. Scheme of the structured illumination microscopy (openSIM) setup** - The light-path (green) shown in the schematic above starts with a laser-module ($\lambda_c = 532\,nm$) which gets expanded by a small beam-expander based on a cellphone- and a biconvex lens by a factor of $\approx 4$. The collimated beam illuminates the Raspberry Pi controlled DMD before it gets relayed by a telescope. In the Fourier-plane of the first lens, a Fourier-filter which blocks the zeroth order. A second mirror followed by a tube-lens creates the diffraction orders resulting from the grating in the BFP of the objective lens. The resulting grating illuminates a sample movable by the z-stage. The detection path (green) follows a typical infinity-corrected microscope where either a CMOS (e.g. IDS, BASLER) or cellphone-camera combined with an eye-piece.

All files to replicate this experiment can be found here.

#### Quantitative Imaging using the (*open*KOEHLER)

An alternative to incoherent imaging methods, where fluorescently labelled cells are captured, is given by quantitative phase-imaging (QPI). This modality is very attractive for biological samples, because it is a label-free method thus omitting the time consuming labelling procedures which are sometimes also altering the behaviour or appearance of the subject of observation. Based on the adaptive illumination scheme described in [3], we incorporated a low-cost HDMI video-projector (40 Euro, Generic brand, China) which adds a fully adaptive illumination source to the system (see Fig. 21). The LCD-panel inside the projector is hold in a customized UC2 module which includes the high-power LED for the illumination, the controlling PCB which translates the incoming HDMI video-signal for the $320 \times 240$ RGB 2,4 inch TFT screen (ILI9341, China) and a set of lenses to ensure correct Koehler illumination.
The LCD-plane is a conjugated plane of the BFP of the microscopic objective lens and can create different illumination schemes like oblique illumination, (quantitative) differential phase contrast ($q$ DPC), Fourier Ptychography Microscopy, Dark-field, etc. by addressing a specific bitmap pattern on the 2D plane. Each pixel produces a plane-wave in the sample-plane if it is in the "on"-state and can transfer certain frequencies of the object. The super-position of all illuminating plane-waves in the camera-plane gives the later image following the "Abbe"-method (see. [5]).

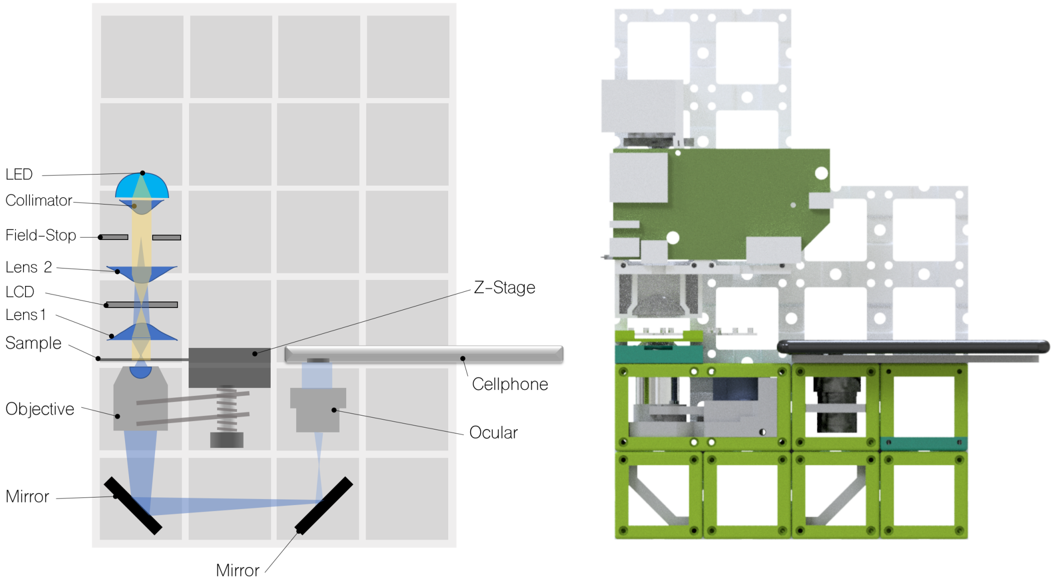

**Fig 21. Ready-to-print *open*KOEHLER module** - The *open*KOEHLER module accommodates a LCD in place of the effective illumination source (i.e. optically conjugate to the BFP) to create an adaptive illumination source (e.g. Koehler-illumination). The module can be controlled using a standard computer equipped with an HDMI port. By varying the pattern in the LCD-plane the contrast of transparent cells captured by a cellphone camera can be optimized.

**Optical Setup** The optical system, shown Fig. 21 follows a Koehler illumination [5], where the condenser aperture plane is imaged into the BFP of the detecting objective lens and a field-stop is imaged into the sample plane to block possible stray-light. Taking the high-power

white LED equipped with a collimating lens and adding two injection-molded aspherical lenses (Thorlabs ACL3026U, $f' = 26\,mm$, $NA = 0.55$) $L_1$ and $L_2$ images the LCD-plane representing the aperture plane into the BFP of the objective lens and further images a field-stop in the sample plane.
A variation of the pattern displayed on the LCD controlled as a secondary display (e.g. HDMI-connection) directly influences the visible contrast. Depending on the displayed pattern (e.g. circle, annulus, etc.), the degree of coherence can be chosen freely.
All files to replicate this experiment can be found here.

#### Educational Areas

The modular concept of building optical setups has proven to be very useful in demonstrations of various principles in microscopy and image formation in general. To exploit this potential, we came up with the idea of *TheBOX*. The comprehensive toolbox provides components for explaining the basics of ray optics, diffraction and different microscopy modalities. It comes in two versions, *Simple* and *Full*. The *SimpleBOX* contains only passive components and covers the optical experiment of secondary and high school level. The *FullBOX* is equipped with electronics like microcontrollers (ESP32) and a Raspberry Pi microcomputer including a 7-inch touch-screen, keyboard and a camera module. In addition to basic experiments, this advanced box can create a compound microscope, a light sheet setup and others which are more suitable for the everyday life in the biological lab. The complete list of setups can be found in the online repository. The target groups for *TheBOX* are schools and other institution that provide courses on optics and microscopy. Thanks to its low price (600 Euro), a school/institution/course organiser can acquire or build multiple boxes and each participant can therefore have access to a hands-on experience, which is nowadays typically not the case. With this we try to become the "Arduino for optics" meaning that the time of getting started is heavily reduced by the plug-and-play nature of the building blocks.
We have successfully tested the concept of *TheBOX* in a series of workshops which are exemplary documented for the Inline Holographic Microscope and the Light sheet hackathon for the "International Day of Light" (IDOL) in the "Lichtwerkstatt Jena" and HHMI Research Institute on building a light sheet setup based on UC2 toolbox from scratch. *TheBOX* has proven to be a useful tool for microscopists, biologists and people generally interested in microscope at all skill-levels to learn e.g. the concept of Fourier optics or study organisms at a cellular level.
A set of trial-runs in Thuringian high schools (Carl-Zeiss-Gymnasium Jena, Montessori Schule Jena, Königin-Luise-Gymnasium Erfurt) to learn how the system can be used for "STEAM"-education (Science, Technology, Engineering, Art and Mathematics) has extremely positive feedback. By conducting interdisciplinary projects, where students study the application of the UC2 system to e.g. track micro-plastic in drinking water using special fluorescent markers. A series of tutorials on how to print, order and assemble can be found here.

**The *"SimpleBOX"*** The *SimpleBOX* is a collection of optical experiments taking place from elementary to high school level. It solely relies on passive optical elements such as lenses, mirrors and objective lenses. The overall material cost is in the range of 150 Euro from known online retailers. The different experiments are listed in the table 1.

**Table 1.** Overview of the experiments inside the *"SimpleBox"*

| **Setup Name** | Description | **Link** |
|---|---|---|
| Keplerian Telescope | Telescope based on two positive lenses to magnify an image by a factor of 2, the image is up-side-down | Github-URL |
| Galilean Telescope | Telescope based on a positive and negative lens to magnify an image by a factor of 2, the image is upright | Github-URL |
| Projector | Projector based on a transparent object (e.g. slide) which gets imaged on a surface using a torch | Github-URL |
| Smartphone microscope | inverted microscope based on an objective lens, two mirrors and an eyepiece | Github-URL |

Each setup comes along with a template which shows how to assemble to optical module as shown in Fig. 3.

**The *"FullBOX"*** [H] *FullBOX* is an extended version of the *SimpleBOX* which adds active components like motors, LEDs and electronics to the cubes making them "smart". Using micro computer like the Raspberry Pi and microcontrollers like the ESP32 or Arduino can create more complex and fully autonomous setups ready for the every-day measurement in the optical lab or for the use in high schools and universities. The overall material cost is in the range of 600 Euro from known online retailers. A list of achievalbe experiments is given by table 1.

**Table 2.** Overview of the experiments inside the *"FullBOX"*

| **Setup Name** | Description | **Link** |
|---|---|---|
| Incubator microscope - transmission | Inverted microscope equipped with an LED array for bright-field microscopic imaging | Github-URL |
| Incubator microscope - epifluorescence | Inverted microscope equipped with a dark-field-like LED illumination for fluorescent microscopic imaging | Github-URL |
| Light sheet microscope | Selective plane imaging using using a Raspberry Pi camera | Github-URL |
| Smartphone microscope | Inverted microscope equipped with an LED array for bright-field microscopic imaging using a cellphone camera | Github-URL |
| Abbe diffraction experiment | Experiment to observe Fourier-filtering by imaging the image and the fourier plane simultaneously | Github-URL |
| Spectrometer | simple spectrometer based on a reflective grating from a CD/DVD | Github-URL |

Using additional components like the DMD-projector based *open*SIM, the *FullBOX* can extended to individual needs.

#### UC2 Use-cases

The core-idea of the UC2-system is to be open, so that it can be used by a large number of people. In best case, users do not only use the system, but participate actively in the iterative design-process by suggesting new applications, finding errors. This can conveniently be done using for example the issue-tracking feature embedded in the GitHub repository. Alternatively private messages, feedback-rounds on workshops or discussions through social media channels such as Twitter can be used as a feedback mechanism. After promoting the principle of the UC2 system in a number of talks and workshops, many people started replicating the system. Since we can not keep track of the number of downloads and actually printed systems, it is hard to track how many people besides us actually build and use it. Nevertheless, we found the scientific community on Twitter, where we created a dedicated UC2-Twitter account (@openUC2)as a helpful measure and feedback mechanism to track issues, ideas, improvements and to give a rough estimate how many systems are in actual use (exemplary shown in Fig. 22).

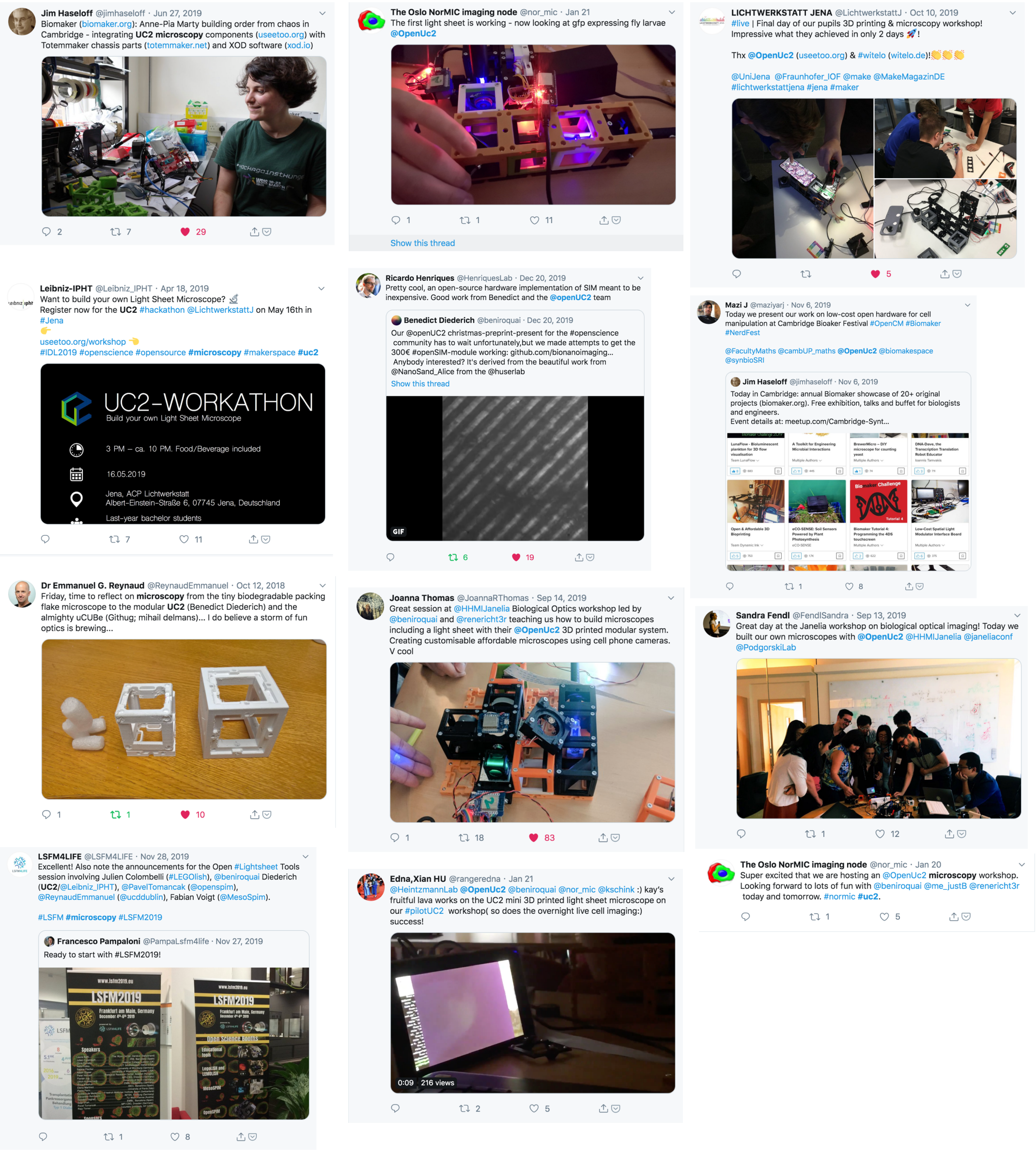

**Fig 22. Publicly announced UC2 workshops and use-cases** *Even though it is hard to track how many UC2 systems are in actual use, we collected an exemplary over-view of some user-feedback and successfully assembled UC2 setups.*

From the workshops we found, that it is of great importance, that the entrance threshold is set very low to attract new users to start developing using the UC2 system. This way, the documentation is aimed to be self-explanatory thus acting as a decentralized multiplication.

#### Sample Preparation

#### Primary Macrophages

Peripheral blood mononuclear cells (PBMCs) were isolated from healthy volunteer adult donors by Ficoll density centrifugation approved by the ethical committee of the university hospital Jena.
Monocytes were detached and $1.5 \cdot 10^5$ were seeded in a $35\,mm$ rinsed with $3\,ml$ X-Vivo 15 with supplements. The blood was mixed with isobuffer (PBS without Ca/Mg (Gibco), $2\,mM$ EDTA (Sigma-Aldrich, St. Louis, USA), 0.1% BSA (Sigma-Aldrich)) and placed on top of Biocoll (Merck, Darmstadt, Germany) without mixing in a $50\,ml$ tube. Biocoll and Blood were centrifuged with 800 x g for $20\,min$ with out break. The ring of PBMCs was transferred in a new $50\,ml$ tube and washed twice with isobuffer. PBMCs were seeded at a density of $1 \cdot 10^6$ $cells/cm^2$ in X-VIVO 15 medium (Lonza, Cologne, Germany) supplemented with 10% (v/v) autologous human serum, 10 ng/ml granulocyte macrophage colony stimulating factor (GM-CSF) and $10\,g/ml$ macrophage colony stimulating factor (M-CSF) (PeproTech, Hamburg, Germany) and Pen/Strep (Sigma-Aldrich). After $1\,h$ PBMCs were washed twice with RPMI and remaining monocytes were then rinsed with X-Vivo with supplements. $16\,h$ after isolation monocytes were washed with prewarmed PBS (w/o Ca/mg) and incubated $7\,min$ with prewarmed with 4 mg/ml lidocain (Sigma-Aldrich) and $1\,mM$ EDTA (Sigma-Aldrich). Detached monocytes were places in a $15\,ml$ tube and centrifuged $7\,min$ by 350 x g. Sediment monocytes were counted and $1.5 \cdot 10^5$ were seeded in a $35\,mm$ dish and rinsed with $3\,ml$ X-Vivo 15 with supplements. After $24\,h$ were the cells washed once with X-Vivo 15 and the monocytes were rinsed with $3\,ml$ fresh X-Vivo 15 with supplements and placed in the microscope.

**Phagocytosis:** Stimulated cells were washed with PBS. X- with/out supplements containing $0.25\,smg/ml$ pHrodo(TM) Green E.coli BioParticles(TM) Conjugate for Phagocytosis (ThermoFisher Scientific, MA, USA) was added and imaged for 1h. Fluorescence intensity was analysed.

#### HPMEC

HPMEC-eGFP were cultured in Endopan 300SL supplemented with serum substitute Panexin SL-S, EGF, FGF-2, VEGF, Vitamin C, R3-IGF-1, GA, Hydrocortisone, Heparin (Pan Biotech, P04-0065K). Cells were maintained in $6\,cm$ culture dishes in $37\,C°/5\%$ CO2 incubator. To prepare the samples, HPMEC-eGFP were passaged at 90 % confluency and 30,000 cells were plated on $12\,mm$ diameter coverslips. After 24 h, cells were fixed in 4 % Paraformaldehyde for $15\,min$ at room temperature and mounted with Mowiol solution.

#### Zebrafish embryo and Drosophila larvae for light sheet imaging

$200\,mg$ Agarose (Agarose standard, art. 3810.2 from Carl Roth GmbH) was dissolved in $10\,ml$ $H_2O$ at $160°C$ while stirring. After Agarose is fully dissolved, temperature is reduced to $100°C$. The tip of a syringe (1 ml syringe, Injekt-F from Braun) was cut before we fill the syringe with ca. $0,25\,ml$ of agarose. The sample was placed (i.e. zebrafish embryo, drosophila larva) inside the agarose using a pipette before covering it with a few drops of liquid agarose using. The agarose needs roughly an hour to solidify in the fridge at around $7°C$.
